# Comparing the evolutionary dynamics of predominant SARS-CoV-2 virus lineages co-circulating in Mexico

**DOI:** 10.1101/2022.07.05.498834

**Authors:** Hugo G. Castelán-Sánchez, Luis Delaye, Rhys P. D. Inward, Simon Dellicour, Bernardo Gutierrez, Natalia Martinez de la Vina, Celia Boukadida, Oliver G Pybus, Guillermo de Anda Jáuregui, Plinio Guzmán, Marisol Garrido Flores, Óscar Fontanelli, Maribel Hernández Rosales, Amilcar Meneses, Gabriela Olmedo-Alvarez, Alfredo Herrera-Estrella, Alejandro Sanchez-Flores, José Esteban Muñoz-Medina, Andreu Comas-García, Bruno Gómez-Gil, Selene Zárate, Blanca Taboada, Susana López, Carlos F. Arias, Moritz U.G. Kraemer, Antonio Lazcano, Marina Escalera-Zamudio

## Abstract

Over 200 different SARS-CoV-2 lineages have been observed in Mexico by November 2021. To investigate lineage replacement dynamics, we applied a phylodynamic approach and explored the evolutionary trajectories of five dominant lineages that circulated during the first year of local transmission. For most lineages, peaks in sampling frequencies coincided with different epidemiological waves of infection in Mexico. Lineages B.1.1.222 and B.1.1.519 exhibited similar dynamics, constituting clades that likely originated in Mexico and persisted for >12 months. Lineages B.1.1.7, P.1 and B.1.617.2 also displayed similar dynamics, characterized by multiple introduction events leading to a few successful extended local transmission chains that persisted for several months. For the largest B.1.617.2 clades, we further explored viral lineage movements across Mexico. Many clades were located within the south region of the country, suggesting that this area played a key role in the spread of SARS-CoV-2 in Mexico.

## INTRODUCTION

Genome sequencing efforts for the surveillance of the Severe Acute Respiratory Syndrome Coronavirus-2 (SARS-CoV-2) has granted public access to a massive number of virus genomes generated worldwide (https://www.gisaid.org/). Exploring SARS-CoV-2 genome data using phylodynamic and molecular evolution tools has allowed researchers to characterize increasing virus diversity ^1^, track emerging viral subpopulations, and explore virus evolution in real-time, both at local and global scales (for examples see ^2–6^). Throughout the development of the COVID-19 pandemic, viral variants have emerged and circulated across different regions of the world ^2,7^, displaying specific mutations that define their phylogenetic patterns ^8,9^. The emergence and spread of SARS-CoV-2 lineages has been routinely monitored since early 2021, informing public health authorities on their responses to the ongoing pandemic ^10^.

Emerging virus lineages are classified using a dynamic nomenclature system (‘Pango system’, Phylogenetic Assignment of Named Global Outbreak Lineages), developed to consistently assign newly generated genomes to existing lineages, and to designate novel virus lineages according to their phylogenetic identity and epidemiological relevance ^11,12^. Virus lineages that may pose an increased risk to global health have been classified as Variants of Interest (VOI), Variants under Monitoring (VUM), and Variants of Concern (VOC), potentially displaying one or more of the following biological properties ^9,13^: increased transmissibility ^14^, decreasing the effectiveness of available diagnostics or therapeutic agents (such as monoclonal antibodies)^15^, and evasion of immune responses (including vaccine-derived immunity) ^10,16,17^. Up to date, five SARS-CoV-2 lineages (including all descending sub-lineages) have been designated as VOC: B.1.1.7 (Alpha), B.1.351 (Beta), P.1 (Gamma), B.1.617.2 (Delta), and B.1.1.529 (Omicron) ^10,18,19^.

Virus lineages that dominate across various geographic regions are likely to have an evolutionary advantage, driven in part by a genetic increase in virus fitness (*i.e*., mutations enhancing transmissibility and/or immune escape) ^20–24^. Moreover, the spread of different VOC across the world has been linked to human movement, often resulting in the replacement of previously dominating virus lineages ^10^. However, exploring lineage replacement and fitness dynamics remains a challenge, as they are impacted by numerous factors, including differential and stochastic growth rates that vary across geographic regions, a shifting immune structure of the host population (linked to viral pre-exposure levels and vaccination rates),^23,24^ and changing social behaviours (such as fluctuating human mobility patterns and the implementation of local non-pharmaceutical interventions across time) ^23,25,26^. Thus, the epidemiological and evolutionary mechanisms enabling some lineages to spread and become dominant across distinct geographic regions, whilst others fail to do so, remain largely understudied.

Mexico has been severely impacted by the COVID-19 pandemic, evidenced by a high number of cumulative deaths relative to other countries in Latin America ^27^. Since the first introductions of the virus in early 2020 and up to November 2021 ^28^, the local epidemiological curve fluctuated between three waves of infection (observed in July 2020, January 2021, and August 2021) ^27–29^. Prior to the first peak of infection, non-pharmaceutical interventions (including social distancing and suspension of non-essential activities) were implemented at a national scale from March 23^rd^ 2020 to May 30^th^ 2020. Nonetheless, a reopening plan for the country was already announced in May 13^th^ 2020, whilst the national vaccination campaign did not begin before December 2020 ^27^. The ‘Mexican Consortium for Genomic Surveillance’ (abbreviated CoViGen-Mex) ^30^ was launched in February 2021, establishing systematic sequencing effort for a genomic epidemiology-based surveillance of SARS-CoV-2 in Mexico. In close collaboration with the national ministry of health, and driven by the sequencing capacity in the country, the program aimed to sequence per month 1,200 representative samples derived from positive cases recorded throughout national territory, based on the proportion of cases reported across states. During May 2021, the sequencing scheme was upgraded to follow the official case report line, in order to better coordinate case reporting and genome sampling across the country.

Over 80,000 SARS-CoV-2 genomes from Mexico are available in GISAID (https://www.gisaid.org/), with approximately one third of these generated by CoViGen-Mex ^30^ (with other national institutions sequencing the rest). From 2020 to 2021 (corresponding to the first year of the epidemic in the country), more than 200 different virus lineages were detected, including all VOC ^19,31^. During this time, different virus lineages co-circulated across national territory, an observation of particular relevance in the context of recombinant SARS-CoV-2 linages that emerged in North America during 2021 ^32^. Some virus lineages also displayed specific dominance and replacement patterns distinct to those observed in neighbouring countries (namely the USA) ^30,33,34^. Taking this into consideration, we hypothesize that the SARS-CoV-2 dominance and replacement patterns observed in Mexico during 2020 to 2021 were driven by lineage-specific mutations impacting local growth rates, further shaped by the immune landscape of the local host population (depending mostly on virus pre-exposure levels at that time). Furthermore, we expect that viral diffusion processes within the country to be associated with local human mobility patterns, and anticipate that the SARS-CoV-2 epidemic in Mexico has been impacted by the epidemiological behaviour within neighbouring countries.

In this light, we set to investigate the introduction, spread and replacement dynamics of five virus lineages that dominated during the first year of the epidemic in Mexico: B.1.1.222, B.1.1.519, B.1.1.7, P.1 and B.1.617.2 ^30,33,34^. For this, we undertook a phylodynamic approach to analyse cumulative SARS-CoV-2 genome data publicly available from Mexico within the context of virus genome data collected worldwide, and further devised a human migration and phylogenetic-informed subsampling approach to increase robustness of tailored phylogeographic analyses. To investigate lineage-specific spatial epidemiology, we contrasted our phylodynamic results to epidemiological and human mobility data from the country, focusing on quantifying lineage importations into Mexico and on characterizing local extended transmission chains across geographic regions (*i.e*., states). Our analysis revealed similar dynamics for the B.1.1.222 and B.1.1.519 lineages, with both likely originating in Mexico, and denoting single extended transmission chains sustained for over a year. For P.1, B.1.1.7 and B.1.617.2 lineages, multiple introduction events were identified, with a few large transmission chains across the country detected. For B.1.617.2 (represented by C5d, the largest and most genetically diverse clade identified), we observed a within-the-country virus diffusion pattern seeding from the south with subsequent movement into the central and north. We further find that Mexico’s southern border may have played an important role in the introduction and spread of SARS-CoV-2 across the country.

## RESULTS

The sampling date of this study comprises January 2020 to November 2021, corresponding to the first year of the epidemic in Mexico, just before the introduction of ‘Omicron’ (B.1.1.529) into the country ^30,33,34^. Our comparative analysis on the temporal distribution of virus lineages in Mexico confirmed previous published observations ^19,33,34^ showing that relative to other virus lineages circulating at the time, only the B.1.1.222, B.1.1.519, B.1.1.7 (Alpha), P.1 (Gamma) and B.1.617.2 (Delta) lineages displayed a dominant prevalence pattern within the country. Moreover, for most of these dominant lineages, peaks in genome sampling frequency (defined here as the proportion of viral genomes assigned to a specific lineage, relative to the proportion of viral genomes assigned to any other virus lineage in a given time point) often coincided with the epidemiological waves of infection recorded (except for B.1.1.7 and P.1) (**Figure 1a and b**).

**Figure 1.**
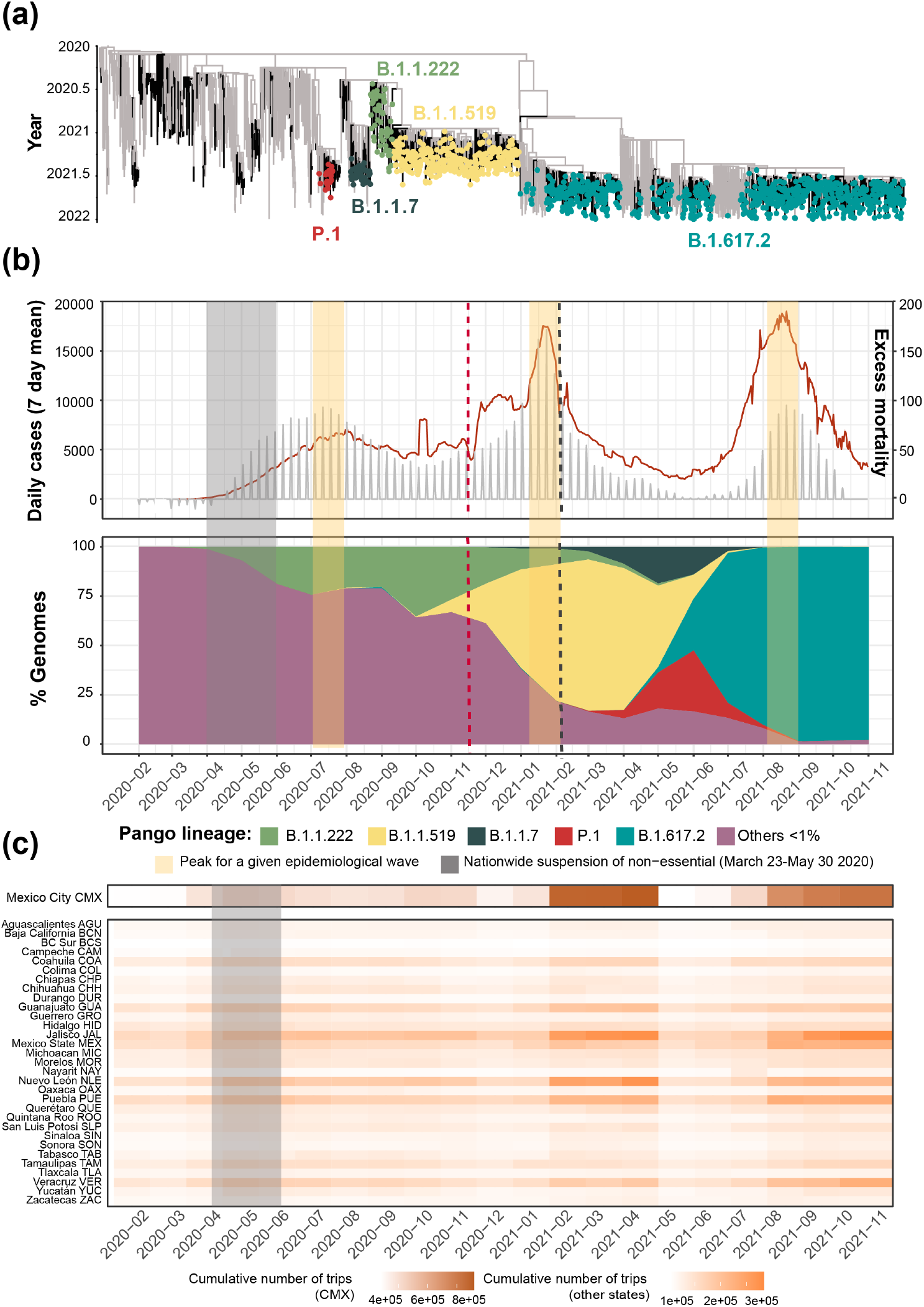
Overview of the SARS-CoV-2 epidemic in Mexico. (a) Time-scaled phylogeny of representative SARS-CoV-2 genomes from Mexico within a global context, highlighting the phylogenetic positioning of B.1.1.222, B.1.1.519, B.1.1.7, P.1 and B.1.617.2 sequences. Lineage B.1.1.222 is shown in light green, B.1.1.519 in yellow, P.1 in red (Gamma), B.1.1.7 (Alpha) in dark green, and B.1.617.2 (Delta) in teal (b) The epidemic curve for COVID-19 in Mexico from January 2020 up to November 2021, showing the average number of daily cases (red line) and associated excess mortality (represented by a punctuated grey curve, denoting weekly average values). The peak of the first (July 2020), the second (January 2021), and the third wave (August 2021) of infection are highlighted in yellow shadowing. The dashed red line corresponds to the start date national vaccination campaign (December 2020), whilst the dashed black line represents the implementation date of a systematic genome sampling and sequencing scheme for the surveillance of SARS-CoV-2 in Mexico (February 2021). The period for the implementation of non-pharmaceutical interventions at national scale is highlighted in grey shadowing. The lower panel represents the genome sampling frequency (defined here as the proportion of viral genomes assigned to a specific lineage, relative to the proportion of viral genomes assigned to any other virus lineage in a given time point) of dominant virus lineages detected in the country during the first year of the epidemic. Lineages displaying a lower sampling frequency are jointly shown in purple. (c) Heatmap displaying the volume of trips into a given state from any other state recorded from January 2020 up to November 2021 derived from anonymized mobile device geolocated and time-stamped data.

During this time, Mexico reported a daily mean test rate ranging between 0.13-0.18 test per 1,000 inhabitants ^29^. Despite a lower testing rate compared to other countries, the cumulative number of viral genomes generated throughout 2020 and 2021 (both by CoViGen-Mex and other national institutions) correlates with the number of cases recorded at a national scale, corresponding to approximately 100 viral genomes per 10,000 cases, or sequencing ~1% of the official COVID-19 cases (**Figure 1 - figure supplement 1**). Although SARS-CoV-2 sequencing remained centralized to Mexico City, the proportion of viral genomes per state also roughly coincided with the spatial distribution of confirmed cases (with Mexico City reporting most cases), as stated officially ^35^ (**Figure 1 - figure supplement 1**). Therefore, SARS-CoV-2 sequencing in Mexico has been sufficient to explore the spatial and temporal frequency of viral lineages across national territory ^30,33,34^, and now to further investigate the number of lineage-specific introduction events, and to characterize the extension and geographic distribution of associated transmission chains, as we present in this study.

### B.1.1.222

The B.1.1.222 lineage circulated in North America between April 2020 and September 2021, mostly within the USA (~ 80% of all B.1.1.222-assigned genomes) and Mexico (~ 20% of all B.1.1.222-assigned genomes). With limited reports from other regions of the world, B.1.1.222 was thus considered as endemic to the region (https://cov-lineages.org/) ^36^. The first B.1.1.222-assigned genome was sampled from Mexico on April 2020 (Mexico/CMX-INER-0026/2020-04-04) ^36^, whilst the last B.1.1.222-assigned genome was sampled from the USA on September 2021 (USA/CA-CDPH-1002006730/2021-09-14). The latest sampling date for B.1.1.222 in Mexico corresponds to July 2021 (Mexico/CHH_INER_IMSS_1674/2021-07-26), two months before the latest sampling date of the lineage at an international scale.

We observe that in Mexico, the B.1.1.222 lineage was continuously detected between April 2020 and May 2021, followed by a steady decline after July 2021 (**Figure 1b**). During its circulation period, most B.1.1.222 genomes were collected from the central region of the country, represented by Mexico City (CMX; **Figure 2a**). For B.1.1.222, a rising genome sampling frequency was observed from May 2020 onwards, coinciding with the first epidemiological wave recorded during July 2020.

**Figure 2.**
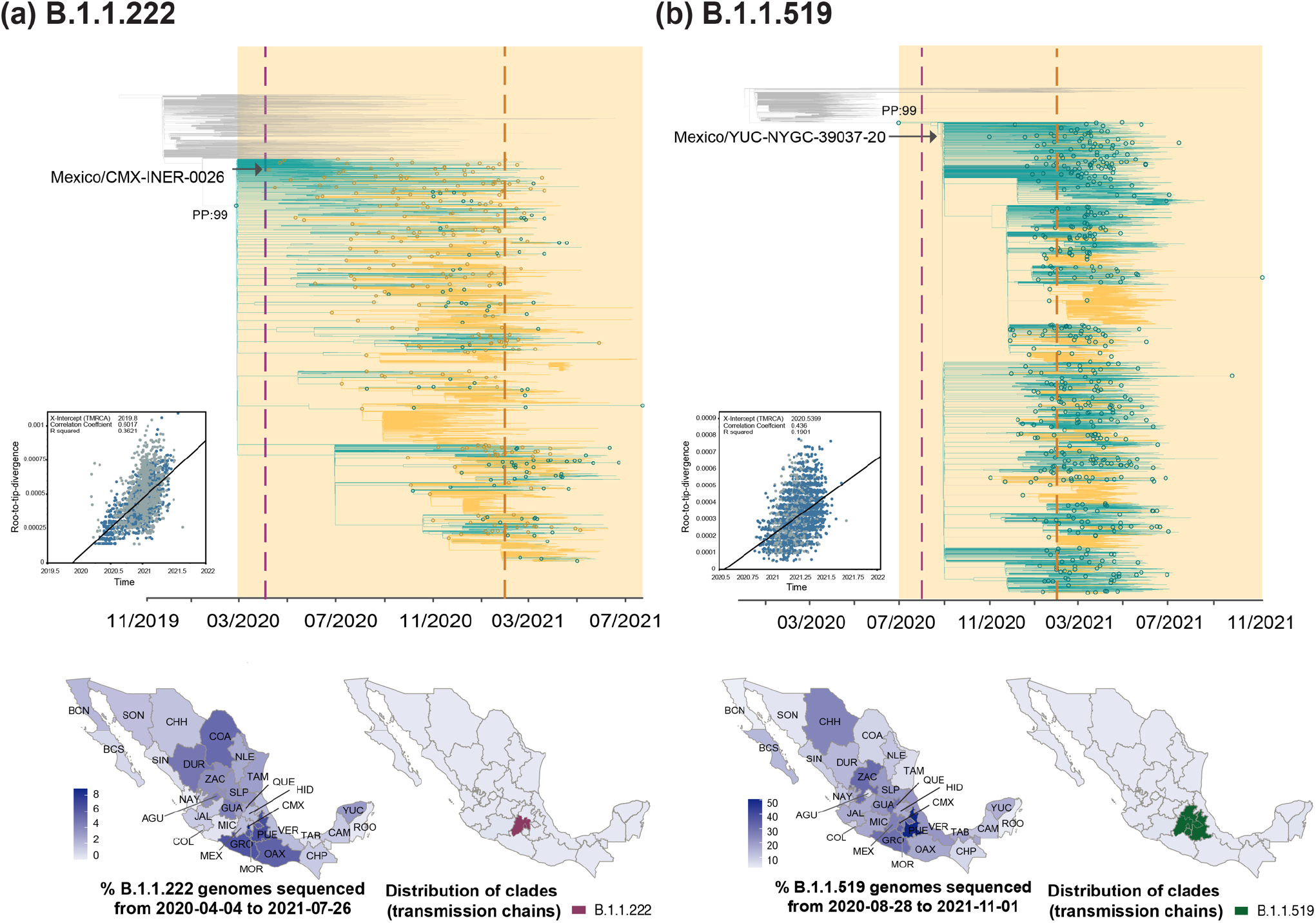
Time-scaled phylogenetic analyses for the B.1.1.222 and B.1.1.519 lineage. Maximum clade credibility (MCC) trees for the (a) B.1.1.222 and (b) B.1.1.519 lineages, in which clades corresponding to distinct introduction events into Mexico are highlighted. Nodes shown as red outline circles correspond to the most recent common ancestor (MRCA) for clades representing independent re-introduction events into Mexico (in teal) or from the USA (in ochre). Based on the earliest and latest MRCAs, the estimated circulation period for each lineage is highlighted in yellow shadowing. The dashed purple line represents the date of the earliest viral genome sampled from Mexico, while its position in the tree indicated. The dashed yellow line represents the implementation date of a systematic virus genome sampling and sequencing scheme for the surveillance of SARS-CoV-2 in Mexico. The corresponding root-to-tip regression plots for each tree are shown, in which genomes sampled from Mexico are shown in blue, whilst genomes sampled elsewhere are shown in grey. Map graphs on the left show the cumulative proportion of genomes sampled across states per lineage of interest, corresponding to the period of circulation of the given lineage (relative to the total number of genomes taken from GISAID, corresponding to raw data before subsampling). Maps on the right represent the geographic distribution of the clades identified.

Subsequently, genome sampling frequency progressively increased to reach a highest of 35% recorded in October 2020, denoting established dominance before the emergence and spread of B.1.1.519 **(Figure 1b)**. Data corresponding to the first year of the epidemic available up to February 2021 (as analysed by Taboada et al. 2021 ^19,33,34^) initially estimated that the B.1.1.222 lineage had reached a maximum genome sampling frequency of approximately 10%. Compared to our results, this revealed an important frequency underestimation (10% vs 35%), attributable to the fact that the vast majority of B.1.1.222-assigned genomes from Mexico (>80%) were generated, assigned and submitted to GISAID after February 2021. Of note, in the USA, the B.1.1.222 lineage reached a maximum genome sampling frequency of 3.5%, compared to 35% observed in Mexico. Thus, more B.1.1.222-assigned genomes from the USA compared to Mexico reflects sequencing disparities across countries ^1^, and contrasts with the region-specific epidemiological scenarios.

Phylodynamic analysis for the B.1.1.222 lineage revealed one main clade deriving from a single MRCA (most recent common ancestor) (**Figure 2a**). The inferred date for this MRCA corresponds to March 2020, denoting a cryptic circulation period of a month (before the earliest sampling date for the lineage within the country, see Methods section 4). The ‘most likely’ location for this earliest MRCA (supported by a relative PP of 0.99) was inferred to be ‘Mexico’, denoting lineage emergence. Thus, subsequent ‘introductions’ should be interpreted as ‘re-introduction’ events into the country (with dates ranging from October 2020 to July 2021). After emergence, B.1.1.222 was seeded into the USA from Mexico multiple times. In this context, we estimate a minimum of 237 introduction events from Mexico into the USA (95% HPD interval = [225-250]), and a minimum of 106 introduction events from the USA into Mexico (95% HPD interval = [95-122]; **Figure 2a**). Based on inferred node dates (for MRCAs) in the MCC tree, the B.1.1.222 lineage displayed a total persistence of up to 16 months.

### B.1.1.519

Directly descending from B.1.1.222 (**Figure 1a**), the B.1.1.519 lineage circulated in North America between August 2020 and November 2021, mostly within the USA (~ 60% of all B.1.1.519-assigned genomes) and Mexico (~ 30% of all B.1.1.519-assigned genomes). As for B.1.1.222, B.1.1.519 genome reporting from other countries was limited, and the B.1.1.519 lineage was also considered as endemic to the region (https://cov-lineages.org/) ^34,37–39^. At an international scale, the earliest B.1.1.519-assigned genome was sampled from the USA on July 2020 (USA/TX-HMH-MCoV-45579/2020-07-31) ^40^, whilst the latest B.1.1.519-assigned genome was sampled from Mexico on December 2021 (Mexico/CHP_IBT_IMSS_5310/2021-12-27) ^41^. During initial phylogenetic assessment, we noted that most of B.1.1.519-assigned genomes collected after November 2021 came from outside North America (namely, from Turkey and Africa). These were further identified as outliers within the tree, likely to be sequencing errors resulting from the use of an inadequate reference sequence for genome assembly, and thus were excluded from further analyses. In Mexico, the B.1.1.519 lineage was first detected on August 2020 (Mexico/YUC-NYGC-39037-20/2020-08-28) ^34^.

Our analysis derived from cumulative genome data from Mexico shows that B.1.1.519 displayed an increasing genome sampling frequency observed between September 2020 and July 2021 (**Figure 1b**). During these months, the spread of B.1.1.519 raised awareness in public health authorities, leading to its designation as a VUM in June 2021 ^10,34,38,39^. During its circulation period, most B.1.1.519 genomes were sampled from the central region of the country, represented by the state of Puebla (PUE; **Figure 2b**). We further observed that by late January 2021, up to 75% of the virus genomes sequenced in Mexico were assigned as B.1.1.519, with the lineage dominating over the second wave of infection recorded (**Figure 1 b**). Similar to B.1.1.222, in the USA, B.1.1.519 only reached a maximum genome sampling frequency of 5% (up to April 2021). Compared to the 75% observed in Mexico, this once again contrast to the epidemiological scenario observed in each country, further exposing sequencing disparities ^36,40^.

Phylodynamic analysis for the B.1.1.519 lineage revealed a similar pattern to the one observed for B. 1.1.222, with one main clade deriving from a single MRCA (**Figure 2b**). The inferred date for this MRCA corresponds to July 2020, again with a ‘most likely’ source location inferred to be ‘Mexico’ (PP: 0.99). Thus, our results suggest that B.1.1.519 circulated cryptically in Mexico for one month prior to its initial detection (**Figure 2b**). After its emergence, the B.1.1.519 lineage was seeded back and forth between the USA and Mexico, with subsequent ‘re-introduction events’ into the country occurring between July 2020 and November 2021. In this light, we estimate a minimum number of 121 introduction events from the USA into Mexico (95% HPD interval = [109-132]), compared to a minimum number of 391 introduction events from Mexico into the USA (95% HPD interval = [380-402]) (**Figure 2b**). Based on inferred node dates in the MCC tree, the B.1.1.519 lineage displayed a total persistence of over 16 months.

### B.1.1.7

The B.1.1.7 lineage was first detected in the UK in September 2020, spreading to more than 175 countries in over a year ^42^. The earliest B.1.1.7-assigned genome from Mexico was sampled on late December 2020 (Mexico/TAM-InDRE-94/2020-12-31), while the latest B.1.1.7-assigned genome was sampled on October 2021 (Mexico/QUE_InDRE_FB47996_S8900/2021-10-13). Our analysis derived from cumulative genome data from the country revealed a continuous detection between February and September 2021. A peak in genome sampling frequency was observed around May 2021, coinciding with a lower number of cases recorded at the time (**Figure 1b**). Our results further confirm that the B.1.1.7 lineage reached an overall lower sampling frequency of up to 25% (relative to other virus lineages circulating in the country), as noted prior to this study (for example, see Zárate et al. 2022) ^27,29,43^. Of interest, similar observations were independently made for other Latin American countries, such as Brazil, Chile, and Peru (https://www.gisaid.org/), likely denoting region-specific dynamics for this lineage.

Phylodynamic analysis for B.1.1.7 revealed an earliest MRCA dating to late October 2020, denoting a cryptic circulation period of approximately two months prior to detection in the country. The earliest genome sampling date also coincides with at least four independent and synchronous introduction events that date back to December 2020 (**Figure 3a**). In total, we estimated a minimum of 224 introduction events into Mexico (95% HPD interval = [219-231]). Potentially linked to the establishment of a systematic genome sequencing in Mexico, most of these were identified after February 2021. Within the MCC, we further identified seven clades (C1a to C7a) representing extended local transmission chains, with C3 and C7 being the largest (**Figure 3a, Supplementary file 2**). During its circulation period, most B.1.1.7 genomes from Mexico were generated from the state of Chihuahua, with these also representing the earliest B.1.1.7-assigned genomes from the country ^34,44^. However, our analysis revealed that only a small proportion of these genomes grouped within a larger clade denoting an extended transmission chain (C2a),with the rest falling within minor clusters, or representing singleton events (**Figure 3a**). Relative to other states, Chihuahua generated an overall lower proportion of viral genomes throughout 2020-2021. Thus, more viral genomes sequenced from a particular state does not necessarily translate into more well-supported clades denoting extended transmission chains, whilst the geographic distribution of clades is somewhat independent to the genome sampling across the country.

**Figure 3.**
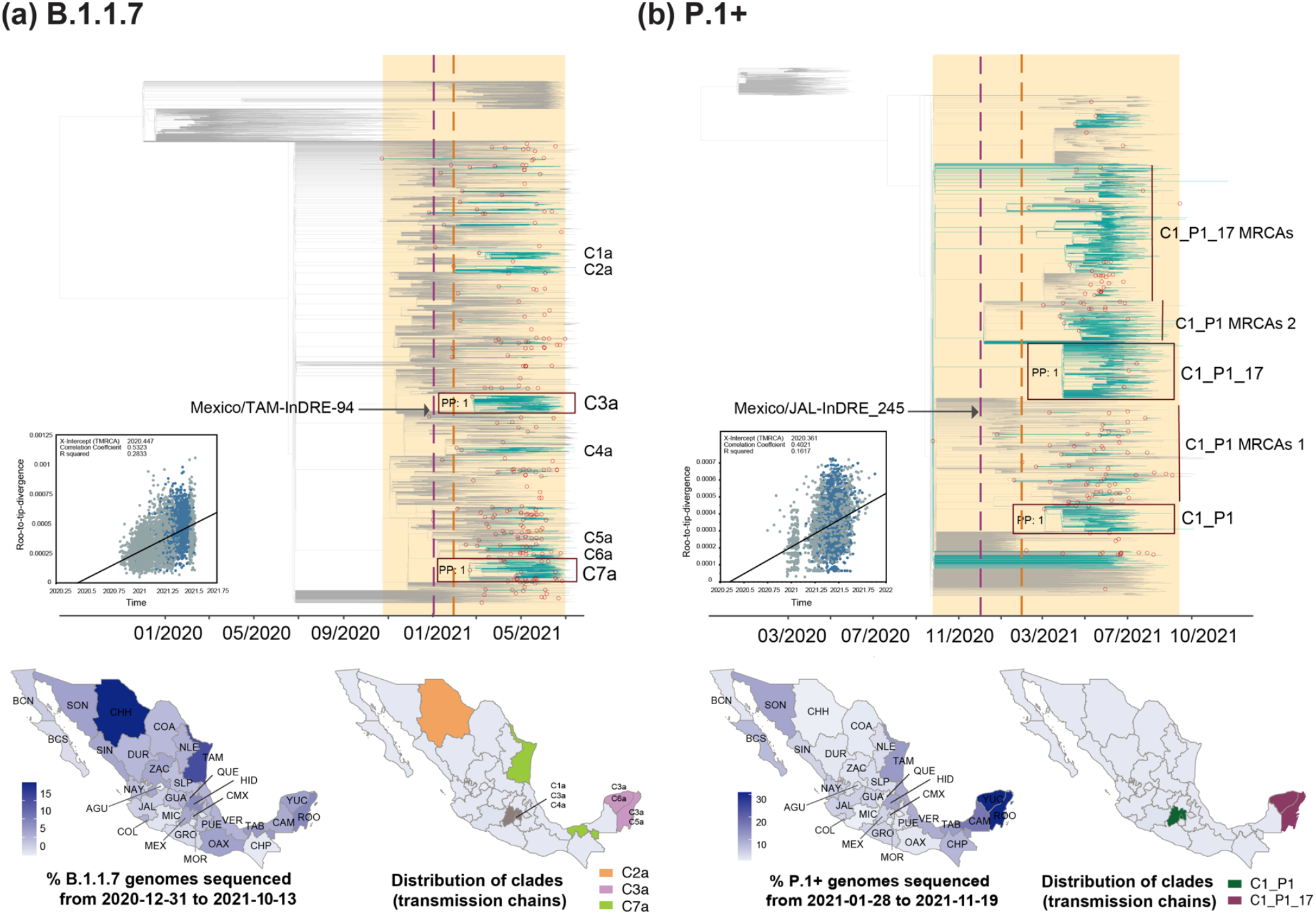
Time-scaled phylogenetic analyses for the B.1.1.7 and P.1 lineages. Maximum clade credibility (MCC) trees for the (a) B.1.1.7 and the (b) P.1 lineages, in which major clades identified as distinct introduction events into Mexico are highlighted. Nodes shown as red outline circles correspond to the most recent common ancestor (MRCA) for clades representing independent introduction events into Mexico. Based on the earliest and latest MRCAs, the estimated circulation period for each lineage is highlighted in yellow shadowing. The dashed purple line represents the date of the earliest viral genome sampled from Mexico, while its position in the tree indicated. The dashed yellow line represents the implementation date of a systematic virus genome sampling and sequencing scheme for the surveillance of SARS-CoV-2 in Mexico. The corresponding root-to-tip regression plots for each tree are shown, in which genomes sampled from Mexico are shown in blue, whilst genomes sampled elsewhere are shown in grey. Map graphs on the left show the cumulative proportion of genomes sampled across states per lineage of interest, corresponding to the period of circulation of the given lineage (relative to the total number of genomes taken from GISAID, corresponding to raw data before subsampling). Maps on the right represent the geographic distribution of the clades identified.

For the larger C3a and C7a clades, both MRCA’s date to February 2021, denoting independent and synchronous introduction events (**Figure 3a**). The C3a comprises genomes collected from 22/32 states in the country, predominantly from Mexico City (CMX), followed by southern states of Yucatán (YUC) and Quintana Roo (ROO) (**Figure 3 - figure supplement 1**). The C3a displayed a persistence of three months: from March to June 2021. For the C7a, viral genomes were sampled from 20/32 states of the country, with >70% of these coming from the southern state of Tabasco (TAB) and north-eastern state of Tamaulipas (TAM) (**Figure 3 - figure supplement 1**). The C7a displayed a persistence of four months: from March to July 2021. Based on inferred node dates within the MCC tree, the B.1.1.7 lineage displayed a total persistence of approximately 10 months.

### P.1

The P.1 lineage was first detected in Brazil during October 2020 ^45^, after which it diverged into >20 sub-lineages that spread to different parts of the world ^19^. Relevant to North America, P.1.17 was the most prevalent sub-lineage detected within the region, again sampled mostly from the USA (~ 60% of all sequences) and from Mexico (~ 30% of all sequences, https://cov-lineages.org/) ^19^. In Mexico, we detected at least 13 P.1 sub-lineages, with the majority of assigned viral genomes belonging to the P.1.17 (66%), and to a lesser extent to the parental P.1 lineage (25%), as was noted prior to this study ^31^. As our dataset comprises viral genomes assigned to the P.1 and descending sub-lineages, it is henceforth referred here as a P.1+.

The earliest P.1+ genome from Mexico was sampled on late January 2021 (Mexico/JAL-InDRE_245/2021-01-28) and the latest on November 2021 (Mexico/ROO_IBT_IMSS_4502/2021-11-19). Cumulative genome data analysis from the country revealed a similar pattern to that observed for B.1.1.7, in which P.1+ genome sampling frequency peaked around April-May 2021, with almost no detection after September 2021. As for B.1.1.7, P.1+ showed an overall lower genome sampling frequency reaching a highest of 25%, again coinciding with a decrease in the number of cases following the second wave of infection recorded (**Figure 1b)**^31,33,34^. During its circulation period in the country, the majority of P.1+ -assigned genomes were sampled from the states Yucatan and Quintana Roo (YUC and ROO; **Figure 3b**).

Our phylodynamic analysis for P.1+ revealed a minimum number of 126 introduction events into Mexico (95% HPD interval = [120-132]). Within the MCC tree, we identified two well-supported clades denoting extended local transmission chains: C1_P1 (corresponding to P.1) and C1_P1_17 (corresponding to P.1.17) (**Figure 3b, Supplementary file 2**). The MRCA of the C1_P1 clade dates to March 2021, showing a persistence of seven months: from March to October 2021. The MRCA of C1_P1_17 dates to October 2020, corresponding to the TMRCA of the global P.1+ clade in the MCC tree. The long branch separating this earliest MRCA and the earliest sampled sequence reveals a considerable lag between lineage emergence and first detection, likely resulting from sub-lineage under-sampling (**Figure 3b**). Therefore, it is not possible to estimate a total lineage persistence based on inferred node dates. Thus, considering tip dates only, the C1_P1_17 clade showed a persistence of five months (earliest collection date: 01/04/2021, latest collection date: 17/09/2021). For the P.1 parental lineage, two clusters of MRCAs representing subsequent introduction events with no evidence of extended transmission were identified (referred here as clade C1_P1 MRCAs 1 and 2). Similarly, for the P.1.17, another cluster of MRCAs representing subsequent introduction events with no evidence of extended transmission was also identified (referred here as C1_P1_17 MRCAs) (**Figure 3b**).

The C1_P1 clade directly descends from viral genomes sampled from South America, and is mostly represented by viral genomes collected from the central region of the country (>40% of these coming from Mexico City and the State of Mexico; CMX and MEX) (**Figure 3 - figure supplement 2**). The C1_P1_17 clade is mostly represented by viral genomes from Mexico (75%), and to a lesser extent by genomes from the USA (20%). ‘Mexico’ genomes are positions basally to the C1_P1_17 clade, collected predominantly from the southern region of the country (>90%, represented by the states of Quintana Roo and Yucatán, ROO and YUC) (**Figure 3 - figure supplement 2**). Overall, our results indicate that in Mexico, the P.1 parental lineage was introduced independently and later than P.1.17, likely from distinct geographic locations. Contrasting to P.1, the P.1.17 lineage displayed a more successful spread, denoted by a sustained transmission chain located to the southern region of the country.

### B.1.617.2

Initially detected in India during October 2020, the B.1.617.2 lineage spread globally to become dominant, and was later associated with an increase in COVID cases recorded globally following March 2021 ^46,47^. The parental B.1.617.2 lineage further diverged into >230 descending sub-lineages (designated as the AY.X) that spread to different regions of the world ^19,46,48^. Again, as our dataset comprises both B.1.617.2 and AY. X-assigned genomes, it is henceforth referred here as a B.1.617.2+.

The first ‘B.1.617.2-like’ genome from Mexico was sampled on September 2020 (Mexico/AGU-InDRE_FB18599_S4467/2020-09-22), followed by a sporadic genome detection throughout January 2021 (with <10 sequences) ^49^. However, the comparative analysis on genome sampling frequencies revealed that expansion of B.1.617.2+ only occurred after April 2021 (**Figure 1b**). We further confirmed that by August 2021, the lineage had reached a relative frequency of >95%, coinciding with the peak of the third wave of infection recorded in the country ^50^. Up to the sampling date of this study, we detected >80 B.1.617.2 sub-lineages (AY.X) circulating in Mexico, with most viral genomes assigned as AY.20 (22%), AY.26 (13%), and AY.100 (5%), followed by AY.113, AY.62 and AY.3. Of interest, these were previously noted to be mostly prevalent within North America (https://cov-lineages.org/) ^51–55^. During its circulation period, B.1.617.2+ displayed a more homogeneous genome sampling distribution across Mexico, as compared to other virus lineages. Again, this is likely to be associated with the establishment of a systematic viral genome sampling and sequencing following February 2021, further driven by the widespread expansion of the lineage throughout the country (**Figure 4**).

**Figure 4.**
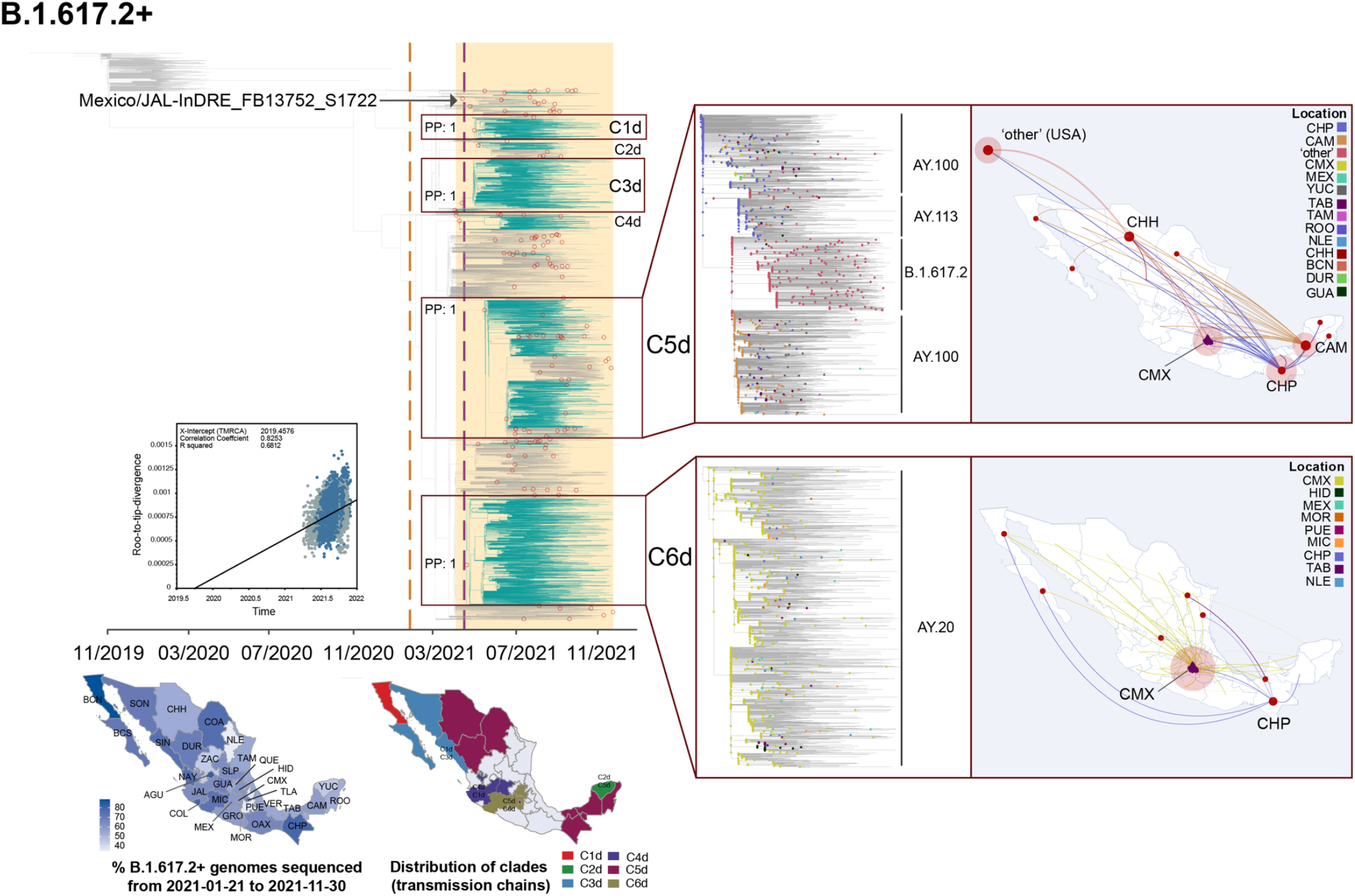
Time-scaled and phylogeographic analysis for the B.1.617.2 lineage. Maximum clade credibility (MCC) tree for the B.1.617.2 lineage, in which major clades identified as distinct introduction events into Mexico are highlighted. Nodes shown as red outline circles correspond to the most recent common ancestor (MRCA) for clades representing independent introduction events into Mexico. Based on the earliest and latest MRCAs, the estimated circulation period for each lineage is highlighted in yellow shadowing. The dashed purple line represents the date of the earliest viral genome sampled from Mexico, while its position in the tree is indicated. The dashed yellow line represents the implementation date of a systematic virus genome sampling and sequencing scheme for the surveillance of SARS-CoV-2 in Mexico. The corresponding root-to-tip regression plot for the tree is shown, in which genomes sampled from Mexico are shown in blue, whilst genomes sampled elsewhere are shown in grey. The map graph on the left show the cumulative proportion of genomes sampled across states per lineage of interest, corresponding to the period of circulation of the given lineage (relative to the total number of genomes taken from GISAID, corresponding to raw data before subsampling). The map on the right represents the geographic distribution of the main clades identified (for further details see Supplementary file 2). On the right, a zoom-in to the C5d and C6d clades showing sub-lineage composition with the most likely location estimated for each node. Geographic spread across Mexico inferred for these clades is further represented on the maps on the right, derived from a discrete phylogeographic analysis (DTA, see Methods section 4). Viral transitions between Mexican states are represented by curved lines coloured according to sampling location, showing only well-supported transitions (Bayes Factor >100 and a PP >0.9) (see Table 1).

Phylodynamic analysis for B.1.617.2+ revealed a minimum number of 142 introduction events into Mexico (95% HPD interval = [125-148]). Within the MCC, six major clades denoting extended transmission chains were identified (C1d-C6d), with C1d, C3d, C5d and C6d being the largest (**Figure 4, Supplementary file 2**). At least four independent introduction events were detected as the earliest (and synchronous) MRCAs, all dating to April 2021 (including the ancestral nodes of the C3d, C4d and C6d clades). Based on inferred node dates in the MCC tree, we report a total lineage persistence of seven months (up to November 30^th^ 2021). C2d comprises ‘Mexico’ virus genomes assigned as AY.62, sampled mainly from the state of Yucatán. Clade C4d comprises genomes from Mexico assigned as AY.3, sampled mostly from the central and south of the country (JAL) (**Figure 4**). Of interest, the C1d and C3d clades represent two independent introduction and spread events of the AY.26 sub-lineage into the country. C1d comprises genomes from Mexico sampled from the north (>60%; BCS, SIN, JAL), followed by central (CMX) and south-eastern states (VER, ROO and YUC) (**Figure 4**). The MRCA of the C1d dates to May 2021, denoting a clade persistence of six months (from May 2021 to November 2021). Comparably, the C3d comprises genomes from Mexico mostly sampled from the north (37%; SIN, BCS and SON). Comparably, the MRCA of the C3d dates to April 2021, denoting a clade persistence of seven months (from April 2021 to November 2021) (**Figure 4**).

For the largest clades identified, C5d comprises viral genomes assigned as AY.100 (44%), to the parental B.1.617.2 (40%), and to the AY.113 (12%). Within this clade, we observe that the AY.100 and B.1.617.2 genomes are separated by a central sub-cluster of AY.113-assigned genomes (**Figure 4**). Approximately 70% of the genome sequences within C5d were sampled from Mexico (mostly assigned as AY.100 and AY.113), whilst 30% were sampled from the USA (mostly assigned as B.1.617.2). The majority of the ‘Mexico’ genomes are positioned basally and distally within the clade, sampled from all 32 states, but predominantly from north, centre and southern regions (>50%; represented by CHH, DUR, NLE, CMX, MEX, CAM, YUC, TAB, CHP and ROO) (**Figure 4**). Thus, the C5d represents the most genetically diverse and geographically widespread clade identified in Mexico. The MRCA of the C5d dates to May, denoting a clade persistence of up to six months (from May 2021 to November 2021). C6d is the second largest clade identified, comprising viral genomes from Mexico assigned as AY.20, mostly collected from central region of the country (>60%; represented by CMX, MEX, MOR, MIC and HID) (**Figure 4**). Thus, contrasting to C5d, C6d denotes an extended transmission chain with a geographic distribution mainly restricted to central Mexico. The MRCA of the C6d clade dates to April, displaying a clade persistence of seven months (from April 2021 to November 2021).

### Spread of B.1.617.2

Given the size and diversity of the C5d and C6 clades, we further explored viral diffusion patterns across the country using a phylogeographic approach (see Methods section 4). For the C5d clade, viral spread is likely to have occurred from the south (represented by the states of Chiapas and Campeche; CHP and CAM) into the rest of the country (**Figure 4 - video 1**). Well-supported transitions (scored under a BF>100 and a PP>.90) were mostly inferred from the southern state of Campeche (CAM) into central and northern states, and subsequently from the northern state of Chihuahua (CHH) into the central and northern region of the country, with some bidirectionality observed. Well-supported transitions were also observed from Baja California into Chihuahua (BCN/BCS into CHH), and from Chihuahua into the USA (arbitrarily represented by the geographic coordinates of the state of California) (**Figure 4**). Contrastingly, for C6d, a limited viral spread was observed from central states (represented by Mexico City, CMX) into central, northern and southern regions of the country (again with some bidirectionality observed). Well-supported transitions were also inferred from the southern state of Chiapas into central and northern region of the country (**Figure 4 - video 2**). Bayes Factor (BF) and Posterior Probability (PP) for well-supported transitions observed between locations can be found as **Table 1**.

**Table 1.** Bayes Factor (BF) and Posterior Probability (PP) for well-supported transitions observed between locations*

| **C5d** |  |  |  | **C6d** |  |  |  |
| --- | --- | --- | --- | --- | --- | --- | --- |
| **Location** |  |  |  | **Location** |  |  |  |
| **From** | **To** | **BFR** | **PP** | **From** | **To** | **BFR** | **PP** |
| BCN | CHH | 14535.32494 | 1 | AGU | CHP | 13635.15617 | 1 |
| CAM | CHP | 14535.32494 | 1 | BCN | CHP | 13635.15617 | 1 |
| CAM | CMX | 14535.32494 | 1 | CHP | CMX | 13635.15617 | 1 |
| CAM | MEX | 14535.32494 | 1 | CHP | COA | 13635.15617 | 1 |
| CAM | MIC | 14535.32494 | 1 | CHP | DUR | 13635.15617 | 1 |
| CAM | other | 14535.32494 | 1 | CHP | GRO | 13635.15617 | 1 |
| CAM | QUE | 14535.32494 | 1 | CHP | GUA | 13635.15617 | 1 |
| CAM | ROO | 14535.32494 | 1 | CHP | HID | 13635.15617 | 1 |
| CAM | SLP | 14535.32494 | 1 | CHP | JAL | 13635.15617 | 1 |
| CAM | SON | 14535.32494 | 1 | CHP | MEX | 13635.15617 | 1 |
| CAM | TAB | 14535.32494 | 1 | CHP | MIC | 13635.15617 | 1 |
| CAM | TAM | 14535.32494 | 1 | CHP | NLE | 13635.15617 | 1 |
| CAM | TLA | 14535.32494 | 1 | CHP | OAX | 13635.15617 | 1 |
| CAM | VER | 14535.32494 | 1 | CHP | other | 13635.15617 | 1 |
| CAM | ZAC | 14535.32494 | 1 | CHP | PUE | 13635.15617 | 1 |
| CMX | CHH | 14535.32494 | 1 | CHP | QUE | 13635.15617 | 1 |
| CHH | CHP | 14535.32494 | 1 | CHP | SIN | 13635.15617 | 1 |
| CHH | CMX | 14535.32494 | 1 | CHP | SLP | 13635.15617 | 1 |
| CHH | DUR | 14535.32494 | 1 | CHP | SON | 13635.15617 | 1 |
| CHH | GUA | 14535.32494 | 1 | CHP | TAB | 13635.15617 | 1 |
| CHH | MIC | 14535.32494 | 1 | CHP | TLA | 13635.15617 | 1 |
| CHH | NLE | 14535.32494 | 1 | CHP | VER | 13635.15617 | 1 |
| CHH | QUE | 14535.32494 | 1 | CAM | CHP | 13635.15617 | 0.998890122 |
| CHH | TAB | 14535.32494 | 1 | NLE | TAB | 13635.15617 | 0.998890122 |
| CHH | TAM | 14535.32494 | 1 | CHP | TAM | 6810.002999 | 0.997780244 |
| CHH | VER | 14535.32494 | 1 | CHP | YUC | 2714.911095 | 0.99445061 |
| CHH | ZAC | 14535.32494 | 1 | MEX | PUE | 164.4591205 | 0.915649279 |
| CAM | CMX | 14535.32494 | 0.998890122 |  |  |  |  |
| CHH | TLA | 14535.32494 | 0.998890122 |  |  |  |  |
| CAM | SIN | 3621.718465 | 0.995560488 |  |  |  |  |
| BCS | CHH | 1023.240732 | 0.984461709 |  |  |  |  |
| MIC | YUC | 468.8988157 | 0.966703663 |  |  |  |  |
| CHH | other | 399.6060762 | 0.961154273 |  |  |  |  |
| CAM | COA | 188.7999953 | 0.921198668 |  |  |  |  |
| MEX | YUC | 126.5111615 | 0.886792453 |  |  |  |  |

\*derived from the phylogeographic analyses for C5d and C6d (B.1.617.2+). Only values of BF>100 and PP>.9 are shown.

### Linking virus spread to human mobility data

Our analysis on human mobility data derived from mobile phone usage (collected between January 2020 and December 2021 at a national scale, see Methods section 5), revealed two mobility peaks across time (**Figure 1c**). The first occurred between February and April 2021, coinciding with the introduction and spread of the B.1.1.7 and P.1+ lineages, and with the contraction of B.1.1.519. The second mobility peak was observed between August and November 2021, coinciding with the expansion of the B.1.617.2 lineage. Increased human movement (represented by the cumulative number of trips into a given state) were observed for Mexico City, and to a lesser extent, for Jalisco, the State of Mexico, Nuevo León and Puebla (followed by Coahuila, Guanajuato and Veracruz) (**Figure 1c**). Mean connectivity within national territory revealed intensified movements from Mexico City into the State of Mexico, Morelos, Hidalgo, Puebla, Veracruz and Jalisco, and from Jalisco into Michoacán and Guadalajara (**Figure 1 - figure supplement 2**). However, for both the C5d and C6d clades, no correlation between viral transitions and mean connectivity was observed (C5d: Adjusted R-squared: 0.006577, F-statistic: 7.15 on 1 and 928 DF, p-value: 0.007628. C6d: Adjusted R-squared: 0.3216, F-statistic: 470.8 on 1 and 990 DF, p-value: < 2.2e-16), nor with the pairwise distance between states (C5d: Adjusted R-squared: 0.01086, F-statistic: 4.051 on 1 and 277 DF, p-value: 0.04511. C6d: Adjusted R-squared: 0.02296, F-statistic: 2.715 on 1 and 72 DF, p-value: 0.1038). Many of the lineage-specific clades we identify displayed a geographic distribution within southern region of the country (*i.e*., clades C3a and C7 for B.1.1.7, clade C1_P1_17 for P.1, and clades C2d and C5d for B.1.617.2). In this context, ranking connectivity between the southern region of the country (represented by Yucatán, Quintana Roo, Chiapas and Campeche) and the remaining 28 states did reveal a consistently high number of bidirectional movements between regions (represented by the CAM, CMX and VER) (**Figure 1 - figure supplement 2**).

## DISCUSSION

Our results reveal contrasting epidemiological and evolutionary dynamics between virus lineages circulating in Mexico during the first year of the epidemic, with some identifiable patterns. Both the B.1.1.222 and B.1.1.519 lineages likely originated in Mexico, characterized by single clades denoting extended sustained transmission for over a year. During this time, both lineages dominated in Mexico, and were seeded back and forth between Mexico and the USA, but never dominated across the USA. Thus, the number of publicly available viral genomes from each country reflects sequencing disparities that contrast with the lineage-specific epidemiological patterns observed across regions, highlighting the need to leverage genomic surveillance efforts across neighbouring nations using joint strategies ^1^.

Further similarities were observed for the P.1 and B.1.1.7 lineages, for which peaks in genome sampling frequencies coincided with a decrease in cases following the second wave of infection. We further confirm that P.1 and B.1.1.7 did not dominate in Mexico ^34,44^, in contrast to what was observed in other countries such as the UK and USA ^23,45,56^. Similar observations were independently made for other Latin American countries (some with better genome representation than others, like Brazil ^45^), suggesting that the overall epidemiological dynamics of B.1.1.7 in Latin America may have differed substantially from that observed in the USA and UK ^23,45,56^. Such differences could be explained partly by competition between lineages, exemplified in Mexico by the regional co-circulation of B.1.1.7, P.1 and B.1.1.519. Nonetheless, a lack of representative number of viral genomes for most of these countries prevents exploring such hypothesis at a larger scale, and further highlights the need to strengthen genomic epidemiology-based surveillance. However, the overall evolutionary dynamics observed for these VOC are comparable to those reported in other countries ^23,45,56^. As an example, in the USA, the earliest introductions reported for the B.1.1.7 lineage were synchronous to those observed in Mexico (occurring between October and November 2020), and were also characterized by a few extended transmissions chains with a distribution often constrained to specific states ^56^. Thus, as drawn from our study, the successful spread of a given virus lineage does not seem to be linked to a higher number of introduction events, but rather to the extent and distribution of transmission chains, with size likely reflecting virus transmissibility ^57^.

In Mexico, the introduction and spread of the B.1.1.7, P.1 and B.1.617.2 lineages was characterized by multiple introduction events, resulting in a few successful extended local transmission chains. The epidemiological and evolutionary dynamics of the three VOC show that not only did these coincide temporally, but also revealed multiple and independent transmission chains corresponding to different lineages (and sub-lineages) spreading across the same geographic regions. Our results further revealed several clades belonging to different virus lineages distributed within the south region of the country, suggesting that this area has played a key role in the spread of SARS-CoV-2. Of notice, such pattern is comparable to what has been observed for arboviral epidemics in Mexico (Gutierrez et al, *in preparation*,^58^). Jointly, such observations indicate that the south region of Mexico (represented by the states of Chiapas, Yucatán and Quintana Roo) may be a common virus entry and seeding point, emphasising the need for an enhanced virus surveillance in these states that share borders with neighbouring countries in Central America, further highlighting the importance of devising tailored surveillance strategies applied to specific states (*i.e*., sub region-specific surveillance).

In general, an increasing growth rate (*Rt*; defined as the instantaneous reproductive number that measures how an infection multiplies ^59^) observed for different SARS-CoV-2 lineages dominating across specific regions can be partially explained by a fluctuating virus genetic background (*i.e*., emerging mutations that impact viral fitness) ^23,45,56^. In light of our results, relative to the parental B.1.1.222 lineage, B.1.1.519 displayed only two amino acid changes within the Spike protein: T478K and P681H ^39^. Mutation T478K locates within the Receptor Binding Domain (RBD), with a potential impact in antibody-mediated neutralization ^60,61^. On the other hand, mutation P681H is located upstream the furin cleavage site, and falls within an epitope signal hotspot ^62^. Thus, it may enhance virus entry ^63^, reduce antibody-mediated recognition ^62^, and confer Type I interferon resistance ^64^. We speculate that at least one of these mutations may have contributed to the dominance of B.1.1.519 over B.1.1.222 by increasing the *Rt*, as has been observed for other SARS-CoV-2 subpopulations ^20,21,65^. In agreement with this observation, in Mexico, an *Rt* of 2.9 was estimated for B.1.1.519 compared to an *Rt* of 1.93 estimated for B.1.1.222 (both calculated during epidemic week 2020-46, coinciding with the early expansion of B.1.1.519 in the country) ^38^.

Notably, the mutations observed for B.1.1.519 were not exclusive to the lineage, as P681H emerged later and independently in B.1.1.7 (and to a lesser extent in P.1, corresponding to 5% of all sampled genomes) ^66,67^, whilst mutation T478K subsequently appeared in B.1.617.2 ^60,63,64^. Although assessing the impact of emerging mutations on lineage-specific fitness requires experimental validation, data derived from the natural virus population evidences amino acid changes at site 681 of the Spike protein have been predominantly fixed in VOC, with some mutations likely to yield an evolutionary advantage ^60,63,64,68–70^. Thus, we propose that a somewhat ‘shared’ genetic background between the B.1.1.519 and B.1.1.7 lineages (as represented by mutation P681H) may have limited the spread of B.1.1.7 across the country. In this context, our finding suggest that the specific dominance and replacement patterns observed in Mexico were driven (to some extent) by lineage-specific mutations impacting growth rate, with competition between virus lineages at a local scale playing an important role.

Nonetheless, lineage-specific replacement and dominance patterns are likely to be shaped by the immune landscape of the local host population ^71^. In Mexico, relatively widespread and constant exposure levels to genetically similar virus subpopulations for extended periods of time (represented by the B.1.1.222 and B.1.1.519 lineages) may have yielded consistently increasing immunity levels in a somewhat still naïve population (with a nationwide seroprevalence of ~33.5% estimated by December 2020 ^72,73^). As more genetically divergent virus lineages were introduced and began to spread across the country (represented by B.1.1.7 and P.1, and later by B.1.617.2), a shift in the local immune landscape is likely to have occurred, impacted by a viral genetic background prompting (a partial) evasion of the existing immune responses. Supporting this observation, a vaccination rate of above 50% was only reached after December 2021 ^29,74^, suggesting that immunity levels during the first year of the epidemic mostly depended on virus pre-exposure levels.

In the context of human movement related to the spread of SARS-CoV-2 in Mexico, the fluctuating mobility patterns we observed for the country were consistent with a decrease in cases following the second and third waves of infection, likely reflecting changes in the colour-coded system regulating travel restrictions leveraged by the risk of infection ^75^. However, contrasting to our expectations on viral diffusion processes to be associated with local human mobility patterns, the geographic distances and overall human mobility trends observed within Mexico did not correlate with the virus diffusion patterns inferred (represented by B.1.617.2). As geographic distances and human mobility cannot be considered potential predictors of SARS-CoV-2 spread in Mexico, viral diffusion could be explained (to some extent) by human movement across borders. Taking this into consideration, it has been proposed that the spread of SARS-CoV-2 in Mexico is linked to human mobility across USA (for example, see ^44^), as we further evidence in this study by the transmission patterns observed for the B.1.1.222 and the B.1.1.519 lineages at an international scale. However, some of the virus diffusion patterns we observed are also congruent with human migration routes from South and Central America, supporting the notion that SARS-CoV-2 spread in Mexico has been impacted by epidemics within neighbouring regions, and further underlines the need to investigate the potential role of irregular migration on virus spread across geographic regions^76–79^.

Limitations of our study include uncertainty in determining source locations for virus introduction events into the country (for most lineages), restricted by regional genome sampling biases ^80,81^. This is further impacted by *i*) an uneven genome sampling across foreign locations and within the country, and *ii*) by a poor viral genome representation for many countries in Central and South America ^49,82^. Such biases are also likely to affect the viral diffusion reconstructions we present, likely rendering them incomplete. However, as SARS-CoV-2 genome sampling and sequencing in Mexico has been sufficient, we are still able to robustly quantify and characterize lineage-specific transmission chains. It is worth highlighting that a differential proportion of cumulative viral genomes sequenced per state does not necessarily mirror the geographic distribution and extension of the transmission chains identified, but rather represents a fluctuating intensity in virus genome sampling and sequencing through time. Thus, a more homogeneous sampling across the country is unlikely to impact our main findings, but could *i*) help pinpoint additional clades we are currently unable to detect, *ii*) provide further details on the geographic distribution of clades across other regions of the country, and *iii*) deliver a higher resolution for the viral spread reconstructions we present. Overall, our study prompts the need to better understand the impact of land-based migration across national borders, and encourages joint virus surveillance efforts in the Americas.

## METHODS

### 1. Data collation and initial sequence alignments

Global genome datasets assigned to each ‘Pango’^11^ lineage under investigation (B.1.1.222, B.1.1.519, B.1.1.7, P.1 and B.1.617.2) were downloaded with associated metadata from the GISAID platform (https://www.gisaid.org/) as of November 30^th^ 2021 ^49,83^. The total number of sequences retrieved for each virus lineage were the following: B.1.1.222 = 3,461, B.1.1.519 = 19,246, B.1.1.7 = 913,868, P.1 = 87,452, and B.1.617.2 = 2,166,874. These also included all SARS-CoV-2 genomes from Mexico available up the sampling date of this study, generated both by CoViGen-Mex and by other national institutions. Viral genome sequences were quality filtered to be excluded if presenting incomplete collection dates, if >1000 nt shorter than full genome length, and/or if showing >10% of sites coded as ambiguities (including N or X). Individual datasets were further processed using the Nextclade pipeline to filter according to sequence quality ^84^. In addition, a set of the earliest SARS-CoV-2 sequences sampled from late 2019 to early 2020 (including reference sequence Wuhan-Hu-1, GenBank accession ID: MN908947.3), and a set of viral genomes representing an early virus diversity sampled up to 31/05/2020 were added for rooting purposes (https://github.com/BernardoGG/SARS-CoV-2_Genomic_lineages_Ecuador). To generate whole genome alignments, datasets were mapped to the reference sequence Wuhan-Hu-1 (GenBank: MN908947.3) using Minimap2 ^85^. Then, the main viral ORFs (Orf1ab and S) were extracted to generate reduced-length alignments of approximately 25,000 bases long, comprising only the largest and most phylogenetically informative coding genome regions (excluding smaller ORFs, UTRs, and short intergenic sequences).

### 2. Migration data and phylogenetically-informed subsampling

To provide an overview for global introductory events into Mexico as a proxy for dataset reduction, we used openly available data describing anonymized relative human mobility flow into different geographical regions based on mobile data usage ^86,87^ (https://migration-demography-tools.jrc.ec.europa.eu/data-hub/index.html?state=5d6005b30045242cabd750a2). For any given dataset, all ‘non-Mexico’ sequences were sorted according to their location, selecting only the top 5 countries representing the most intense human mobility flow into Mexico. In the case reported sub-lineages, the subsampled datasets were further reduced by selecting the top 5 sub-lineages that circulate(d) in the country. The ‘Mexico’ genome sets were then subsampled to ~4,000 in proportion to the total number of cases reported across time (corresponding to the epidemiological weeks from publicly accessible epidemiological data from the country ^88^). This yielded datasets of a maximum of 8,000 genomes, with an approximate 1:1 ratio of ‘Mexico’ to ‘globally sampled’ viral genomes (keeping those corresponding to the earliest and latest collection dates, sampled both from Mexico and globally). Preliminary Maximum Likelihood (ML) tress were then inferred using IQ-TREE (command line: iqtree - s -m GTR+I+G -alrt 1000) ^89^.

Phylogenetically-informed subsampling is based on maintaining basic clustering patterns, whilst reducing the noise derived from overrepresented sequences. This approach was applied to the ML trees resulting from the abovementioned migration-informed subsampled datasets, by using a modified version of Treemmer v0.3 (https://github.com/fmenardo/Treemmer/releases) to reduce the size and redundancy within the trees with a minimal loss of diversity ^90^. For this, the -lm command was initially used to protect ‘Mexico’ sequences and those added for rooting purposes. During the pruning iterations, the -pp command was used to protect ‘Mexico’ clusters and pairs of ‘non-Mexico’ sequences that are immediately ancestral or directly descending from these. This rendered reduced-size representative datasets that enable local computational analyses. As a note, clades may appear to be smaller relative to the raw counts of genomes publicly available, but actually reflect the sampled viral genetic diversity. Datasets were then used to re-estimate the ML trees, and used an input for time-scaled phylogenetic analysis (see Methods section 4). Our subsampling pipeline is publicly accessible at (https://github.com/rhysinward/Mexico_subsampling).

We further sought to validate our migration-informed genome subsampling scheme (applied to B.1.617.2+, representing the best sampled lineage in Mexico). For this, an independent dataset was built using a different migration sub-sampling approach, comprising all countries represented by B.1.617.2+ sequences deposited in GISAID (available up to November 30^th^ 2021). In order to compare the number of introduction events, the new dataset was analysed independently under a time-scaled DTA (as described in Methods Section 4). The distribution plots for each genome dataset before and after applying our migration- and phylogenetically-informed subsampling pipeline, and a full description of the approach employed to validate our migration-informed subsampling is available as **Appendix 1.**

### 3. Dataset assembly for initial phylogenetic inference

Given the reduced size of the original B.1.1.222 dataset, all sequences retained after initial quality filtering were used for further analyses. This resulted in a 3,849-sequence alignment (including 760 genomes from Mexico). All other datasets (B.1.1.519, B.1.1.7, P.1 and B.1.617.2) were processed under the pipeline described (in Methods section 2) to render informative datasets for phylogeographic analysis. The B.1.1.519 final dataset resulted in a 5,001-sequence alignment, including 2,501 genomes from Mexico. The B.1.1.7 final dataset resulted in a 7,049-sequence alignment, including 1,449 genomes from Mexico. The P.1 final dataset resulted in a 5,493-sequence alignment, including 2,570 genomes from Mexico. The B.1.617.2 final dataset resulted in a 5,994-sequence alignment, including 3,338 genomes from Mexico. All genome sequences used are publicly available and are listed in **Supplementary file 1**. Individual datasets were then used for phylogenetic inference as described above, with the resulting trees inputted for a time-scaled analyses.

### 4. Time-scaled analysis

Output ML trees were assessed for temporal signal using TempEst v1.5.3 ^91^, removing outliers and re-estimating trees when necessary. The resulting trees were then time-calibrated informed by tip sampling dates using TreeTime ^92^ (command line: treetime -aln --tree --clock-rate 8e-4 --dates --keep-polytomies --clock-filter 0). Due to a low temporal signal, a fixed clock rate corresponding to the reported viral evolutionary rate estimated (8× 10^-4^ substitutions per site per year) was used ^93,94^. Root-to-tip regression plots for the ML trees (prior to time calibration, and excluding rooting sequences) show a weak temporal signal, and support the use of a fixed molecular clock rate (8×10^-4^) for the temporal calibration of phylogenetic trees (**Figures 2 - 4**).

To further quantify lineage-specific ‘introduction events’ into Mexico and characterize clades denoting local extended transmission chains, the time-calibrated trees were used as input for a discrete trait analysis (DTA, or ‘discrete phylogeographic inference’), using BEAST v1.10.4 to generate maximum clade credibility (MCC) trees ^95–97^. A DTA approach was suitable for all cases, as only a few discrete locations relatively well sampled across time were considered ^97^. Using fixed ‘time-calibrated’ trees as an input for the DTA is an effective way of circumventing the restrictions of computationally-expensive analyses on large datasets ^95^. Although this approach allows to infer dated introduction events into the study area, it does not consider phylogenetic uncertainty. Thus, the most recent common ancestor ‘MRCA’ dates we report come without credibility intervals. For all introduction events identified, the mean and associated HPD interval were assessed. Following a similar strategy as described in du Plessis et al.^3^ ‘Mexico’ clades were identified as those composed by a minimum of two sister ‘Mexico’ viral genome sequences directly descending from another ‘Mexico’ sequence. Extended local transmission chains were identified as clades composed by >20 viral genome sequences, with at least 80% of these sampled from Mexico, and with ancestral nodes supported by a PP value of >.80. Based on the MCC trees, we further estimated ‘total persistence’ times for the lineages studied, defined as the ‘interval of time elapsed between the first and last inferred introduction events associated with the MRCA of any given clade from Mexico’. On the other hand, the lag between the earliest introduction event (MRCA) and the earliest sampling date for any given lineage corresponds to a ‘cryptic transmission’ period.

For the B.1.1.7, P.1 and B.1.617.2 datasets, analyses were performed to estimate the number of transitions into Mexico from other (unknown) geographic regions. Thus, two locations were considered: ‘Mexico’ and ‘other’. For the B.1.1.222 and B.1.1.519 datasets, we estimated the number of transitions between Mexico and the USA, based on the fact both these lineages were considered endemic to North America (with >90% of the virus genomes sampled from the USA and Mexico) ^36^. For this cases, three distinct geographic locations were considered: ‘Mexico’, ‘USA’ and ‘other’. The ‘most likely’ locations for lineage emergence were further obtained by comparing relative posterior probabilities (PP) between inferred ancestral locations for the given TMRCAs ^95–97^. For all analyses, independent Monte Carlo Markov Chain (MCMC) were run for 10^6^ iterations, sampling every 10^3^ states. To assess for sufficient effective sample size values (*i.e*., ESS>200) associated with the estimated parameters, we inspected MCMC convergence and mixing using Tracer 1.7 ^98^. In the case of B.1.617.2, we further explored viral diffusion patterns across the country by running two additional DTAs applied to the largest monophyletic clades identified within the MCC tree (C5d and C6d). For this, we used 33 distinct sampling locations (including all 32 states from Mexico, plus an ‘other’ location, referring to viral genomes sampled from outside the country). Visualization of the viral diffusion patterns was performed using SpreadViz (https://spreadviz.org/home), an updated web implementation of the Spatial Phylogenetic Reconstruction of Evolutionary Dynamics software SpreaD3 ^99^. In order to identify well-supported transitions between locations ^97^, SpreadViz was also further used to estimate Bayes Factor (BF) values.

### 5. Human mobility data analysis and exploring correlations with genomic data

Human mobility data used for this study derived from anonymized mobile device locations collected between 01/01/2020 and 31/12/2021 within national territory, made available by the company Veraset ^100^. The source dataset includes anonymized identifiers for mobile devices, geographical coordinates (latitude and longitude) and a timestamp. The dataset was used to construct aggregated inter-state mobility networks, where nodes are defined as each of the 32 states from the country, whilst (weighed and directed) edges represent the normalized volume of observed trips between nodes ^100^. The resulting networks were then used to quantify the number of cumulative trips from any state into a given specific state across time, the geographic distances among states, the mean inter-state connectivity observed between April 2021 and November 2021 (corresponding to the expansion period for the B.1.617.2 lineage, see **Figure 4b**, **Supplementary file 3**), and finally, for ranking connectivity between the south region of the country (represented by the states of Yucatán, Quintana Roo, Chiapas and Campeche) and the remaining 28 states (**Supplementary file 3**). The connectivity measure was defined as the sum of the weights for edges that go from any given node into other node(s), reflecting the number of trips in any direction. We then used the ‘PhyCovA’ software tool (https://evolcompvir-kuleuven.shinyapps.io/PhyCovA/) to perform preliminary analysis for exploring the human mobility data from the country as a potential predictor of viral transition among locations ^101^. ‘PhyCovA’ was chosen as an explanatory approach over a fully-integrated GLM implemented in the Bayesian BEAST framework, as the last one would imply a high computational burden related to our datasets ^96^.

## DATA AVAILABILITY

Virus genome IDs and GISAID accession numbers for the sequences used in each dataset are provided in the **Supplementary file 1**. All genomic and epidemiological data supporting the findings of this study is publicly available from GISAID/GenBank, from the Ministry Of Health Mexico^102^, and/or from the ‘Our World in Data’ coronavirus pandemic web portal ^29^. For the GISAID data used, the corresponding acknowledgement table is available on the ‘GISAID Data Acknowledgement Locator’ under the EPI_SET_20220405qd and EPI_SET_20220215at keys ^49^. Our bioinformatic pipeline implementing a migration data and phylogenetically-informed sequence subsampling approach is publicly available at https://github.com/rhysinward/Mexico_subsampling.

## COMPETING INTERESTS

The authors declare no competing interests.

## ACKNOWLEDGEMENTS

HGCS is supported by funding through the “Vigilancia Genómica del Virus SARS-CoV-2 en México” grant from the National Council for Science and Technology-México (CONACyT). SD acknowledges support from the *Fonds National de la Recherche Scientifique* (F.R.S.-FNRS, Belgium; grant n°F.4515.22), from the Research Foundation - Flanders (*Fonds voor Wetenschappelijk Onderzoek-Vlaanderen*, FWO, Belgium; grant n°G098321N), and from the European Union Horizon 2020 project MOOD (grant agreement n°874850). MEZ is currently supported by Leverhulme Trust ECR Fellowship (ECF-2019-542). OGP acknowledges support of the Oxford Martin School. MK and RPDI acknowledge support from the European Union Horizon 2020 project MOOD (#874850). The contents of this publication are the sole responsibility of the authors and do not necessarily reflect the views of the European Commission. The mobility team [MHR, AM, OF, MF, PG, GO, GAJ] and AHE are supported by ‘Fondo Conjunto de Cooperación México-Uruguay’ (Agencia Mexicana de Cooperación Internacional para el Desarrollo). CFA acknowledges support from grants “Vigilancia Genómica del Virus SARS-CoV-2 en México-2022” (PP-F003) from the National Council for Science and Technology-México (CONACyT), grant 057 from the “Ministry of Education, Science, Technology and Innovation (SECTEI) of Mexico City”, and grant “Genomic surveillance for SARS-CoV-2 variants in Mexico” from the AHF Global Public Health Institute at the University of Miami. AL work was supported by DGAPA-PAPIIT (IN214421) and DGAPA-PAPIME (PE204921) of UNAM. We thank Verity Hill, Philippe Lemey, Tim Blokker and Sam Hong for their valuable advice on the technical details related to the methodology used for the time-scaled analyses. We thank all members of the Consorcio Mexicano de Vigilancia Genómica (CoViGen-Mex) for their efforts on sample collection and generating genetic sequence and metadata. Particularly, we thank León Martínez-Castilla and José Campillo Balderas for their contributions in the initial collation of preliminary data. We gratefully acknowledge all data contributors for the GISAID sequence data: *i.e*. the authors and their originating laboratories responsible for obtaining the specimens, and their submitting laboratories for generating the genetic sequence and metadata and sharing via the GISAID initiative, on which this research is based.

## SUPPLEMENT LEGENDS

**Figure 1-figure supplement 1.**
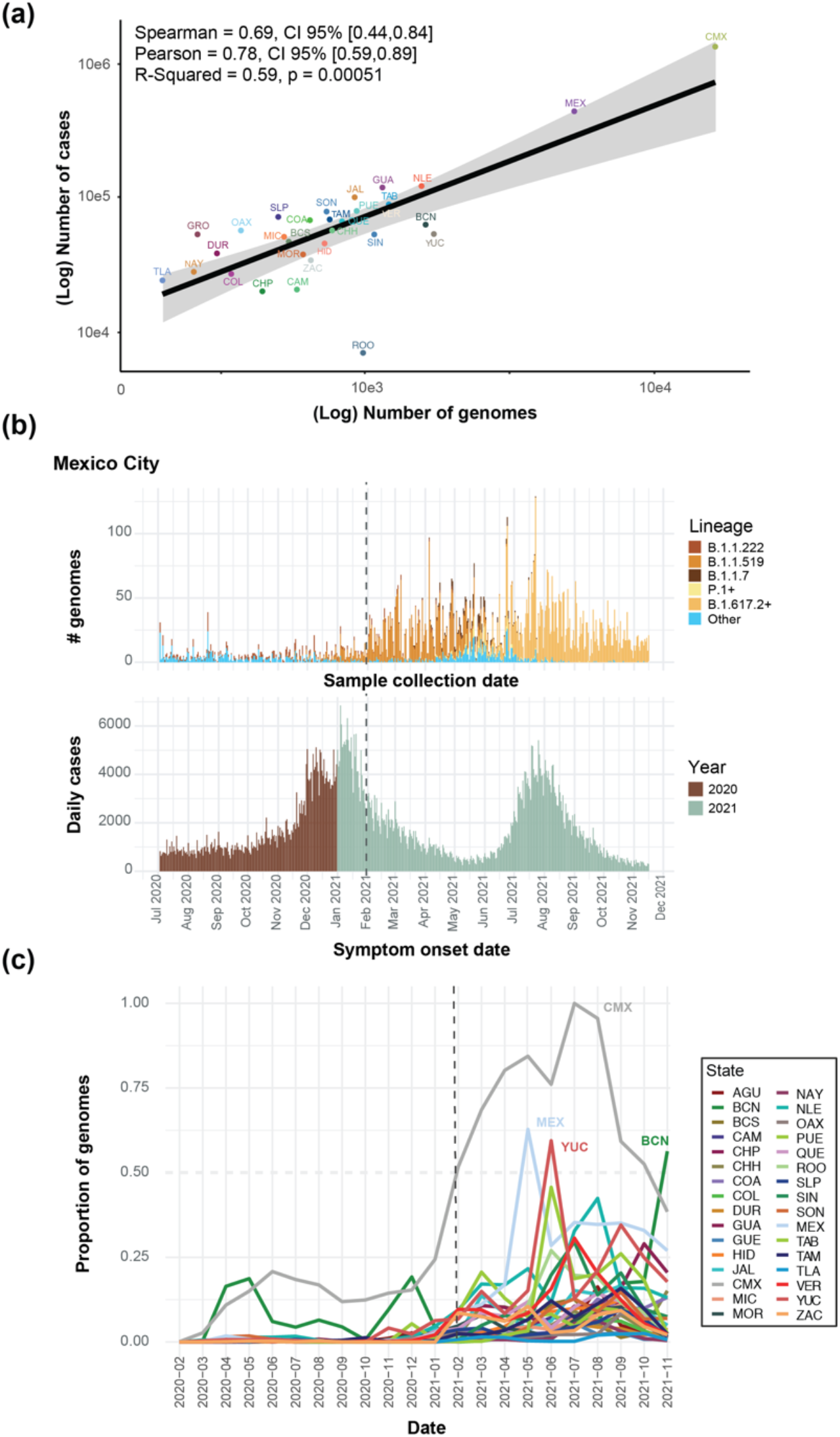
Cumulative number of genome sequences generated per state (data available up to March 2022) (a) A significant correlation between the cumulative number of cases per state versus the number of viral genome sequences available per state is observed, indicating the estimated Spearman/Pearson coefficients and associated 95% confidence intervals (CI). Mexico City (CMX) displays the highest number of genomes sequenced relative to the reported number of cases. (b) A comparison between the total number of genomes sequenced from Mexico City (CMX) assigned to the lineages of interest plotted against collection date, and the number of daily cases reported for Mexico City (CMX) with symptom onset dates ranging from July 2020 up to November 2021 (coloured according to the year of sample collection). The dashed black line represents the implementation date of a broader viral genome sampling and sequencing scheme for the surveillance of SARS-CoV-2 in Mexico (February 2021). (c) The cumulative proportion of genomes sequences generated per state across time (data from February 2020 up to November 2021). The states that generated a proportion of genome sequences above 0.50 (represented by a dashed grey line, relative to other states) are indicated: Mexico City (CMX-grey), State of Mexico (MEX-light blue), Yucatan (YUC-red) and Baja California Norte (BCN-dark green). Once more, the dashed black line represents the implementation date of a broader viral genome sampling and sequencing scheme for the surveillance of SARS-CoV-2 in Mexico (February 2021).

**Figure 1-figure supplement 2.**
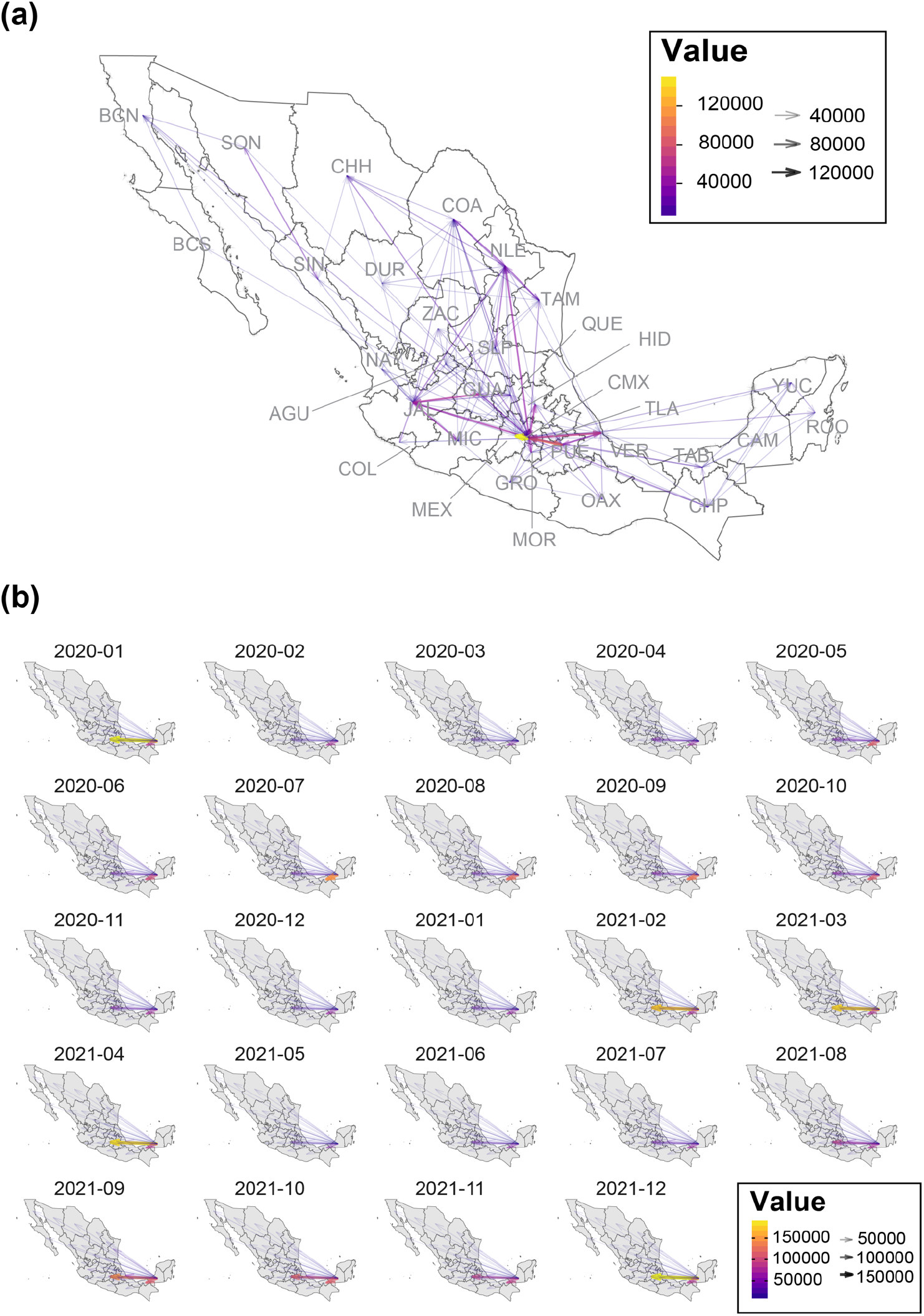
Mean interstate connectivity recorded between 2021 and 2022. (a) Map graph showing the mean intra-state connectivity recorded within national territory, derived from anonymized mobile device locations collected between 01/01/2020 and 31/12/2021. Values above 4E4 are indicated using a colour gradient, whilst arrow thickness within the map represents the total number of bidirectional movements between states. (b) Maps graphs showing the mean inter-state connectivity between the southern region of the country (represented by the states of Yucatán, Quintana Roo, Chiapas and Campeche) and the remaining 28 states (recorded between 01/01/2020 and 31/12/2021). Again, values above 10^-4^ are indicated using a colour gradient, whilst arrow thickness within the map represents the total number of bidirectional movements between states

**Figure 3-figure supplement 1.**
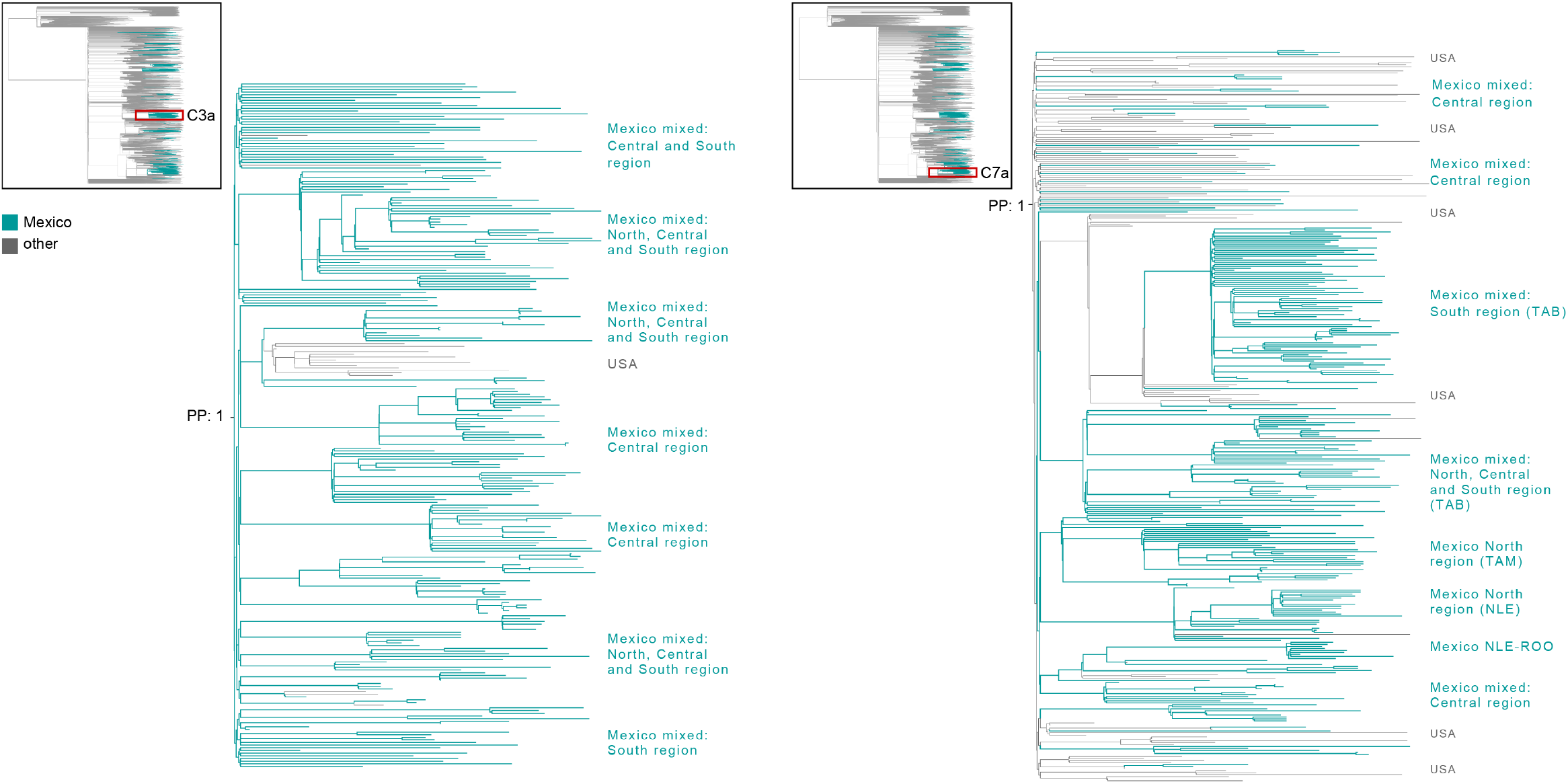
Largest ‘Mexico’ clades within the B.1.1.7 MCC tree. Zoom-in on the C3a and C7a clades identified as the largest within the B.1.1.7 MCC tree. Branch sampling locations are indicated only for sub-clusters composed of >5 sequences. The C3a clade is composed of 254 genome sequences sampled from 22/32 states in the country, mostly from Mexico City (CMX) and State of Mexico (MEX). Clade C7a is composed of 364 genome sequences, sampled mostly from the southern state of Tabasco (TAB). For details of all genome sequences within each clade see Supplementary file 2.

**Figure 3- figure supplement 2.**
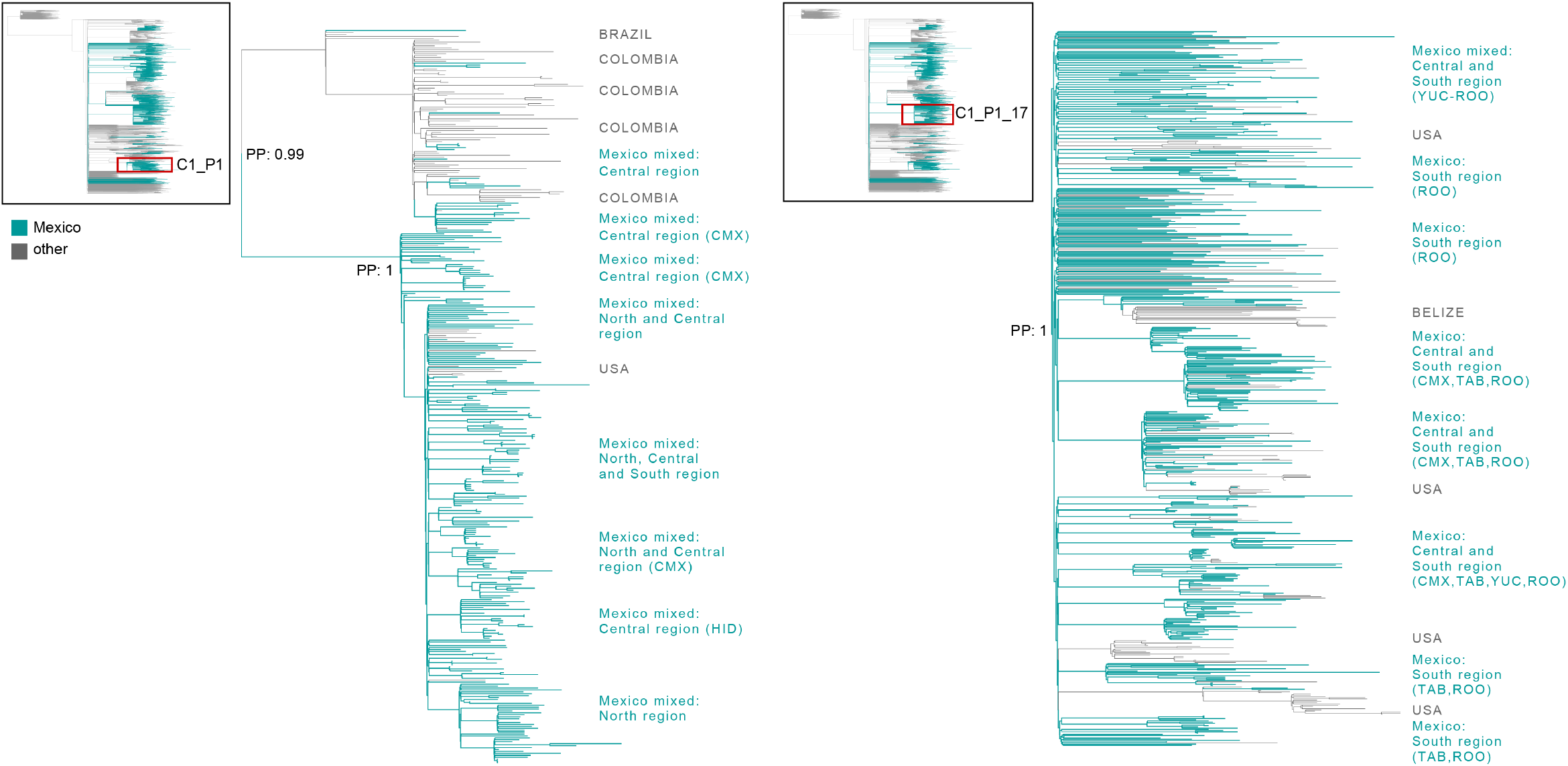
Largest ‘Mexico’ clades within the P.1+ MCC tree. Zoom-in on the C1_P1 and C1_P1_17 clades, identified as the largest within the P.1+ MCC tree. Branch sampling locations are indicated for large sub-clusters composed of >5 sequences. The C1_P1 clade is composed of 277 genome sequences, mostly sampled from the central region of Mexico City (CMX). The C1_P1_17 clade is composed of 588 genome sequences, mostly sampled from the southern states of Quintana Roo (ROO) and Yucatán (YUC). For details of all genome sequences within each clade see Supplementary file 2.

**Video 1.** Animated visualizations of the spread pattern inferred for the C5d clade across Mexico derived from the DTA phylogeographic analysis.

**Video 2**. Animated visualizations of the spread pattern inferred for the C6d clade across Mexico derived from the DTA phylogeographic analysis.

**Supplementary file 1**

Virus genome IDs and GISAID accession numbers for the sequences used in each dataset

**Supplementary file 2**

Full list of names of all genome sequences within each major clade identified for each virus lineage

**Supplementary file 3**

Mobility matrixes summarizing: 1. Ranking connectivity between the southern region of the country, 2. Pairwise distances between states, 3. Mean intrastate connectivity

## Appendix 1 Description of the approach employed to validate our migration-informed subsampling

### METHODS

Our migration-informed approach aims to mitigate the effects of geographical over-representation impacting phylogeographic analyses, in which some regions may appear more frequently as seeding sources than they actually are. Applied to the B.1.617.2 (representing the best sampled lineage in Mexico), we sought to further validate our migration-informed genome subsampling approach by analysing an independent dataset built using a different migration sub-sampling scheme. This new scheme now comprised sub-sampling from all countries represented by B.1.617.2+ sequences deposited in GISAID (available up to November 30^th^ 2021). For this, complete virus genome sequences assigned to the B.1.617.2+ lineage collected globally up to November 30^th^ 2021 available from GISAID (https://www.gisaid.org/) were downloaded as of December 26^th^ 2022. Genome sequences were quality filtered and retained according to the criteria stated in Methods Section 1 (see main text). Genome sequences were further sub-sampled in order to obtain an equal and homogeneous spatial and temporal representation of all geographic regions represented by all the sequences downloaded from GISAID (*i.e*., countries), keeping the number of sampled sequences proportional to the number of cases officially reported from Mexico (corresponding to the epidemiological weeks matching the circulation period of the B.1.617.2 lineage within the country). We further added the set of earliest SARS-CoV-2 sequences sampled globally (‘ROOT’ outgroup), and the 3,320 subsampled genome sequences available from Mexico (used in the original B.1.617.2+ dataset, as described in Methods Section 2, main text). The final dataset resulted in an alignment of 25,107 columns and 6,912 sequences. Subsampling resulted in a homogeneous representation of ≈ 70 countries with an equal number of sequences (10-80 genome per country) per country, relative to their representation in GISAID (**Figure 1**). From these, approximately 3,570 genome sequences were sampled from any other country (*i.e*., ‘global’ sample), preserving the 1:1 sequence ratio of ‘Mexico’ vs ‘non-Mexico’ sequences. The resulting new dataset was further processed and analysed to infer the number of introduction events into Mexico (corresponding to MRCA nodes associated to independent ‘Mexico’ clades), following the steps described in Methods Section 4 (in main text).

### RESULTS AND DISCUSSION

In the new dataset, <100 genome sequences from the USA were retained for further analysis (**Figure 1**), compared to approximately 2,000 genome sequences from the USA included in the original B.1.617.2+ alignment. Thus, we expected a lower number of inferred introduction events into Mexico, as an under-sampling of viral genome sequences from the USA is likely to result in ‘Mexico’ clades not fully segregating (particularly impacting C5d). Our original results revealed a minimum number of 142 introduction events into Mexico (95% HPD interval = [125-148]), with 6 major clades identified (denoting extended transmission chains). The DTA results derived from the new dataset (subsampling all countries) revealed a minimum number of 84 introduction events into Mexico (95% HPD interval = [81-87]), with again 6 major clades identified (**Figure 2**).

Thus, a significantly lower number of introduction events into Mexico were inferred, as was expected. On the other hand, the number of clades identified were consistent between both datasets, supporting for the robustness of our phylogenetic methodological approach. However, in the new dataset, we observe that C5d displayed a reduced diversity, represented by the AY.113 and AY.100 genomes from Mexico, but excluded the B.1.617.2 genome sampled from the USA (as seen in Taboada et al, 2022). This highlights the relevance of our genome sub-sampling using migration data as a proxy. In further agreement with our observations, publicly available data on global human mobility (https://migration-demography-tools.jrc.ec.europa.eu/data-hub/index.html?state=5d6005b30045242cabd750a2) shows that migration into Mexico is mostly represented by movements from the USA, followed by Indonesia, Guatemala, Belize and Colombia. However, the volume of movements from the USA into Mexico is much higher (up to 6 orders of magnitude above the volumes recorded into Mexico from any other country). This further supports for our migration informed subsampling approach of selecting only the top 5 countries with the highest migration rate into Mexico.

#### Appendix 1- Figure legends

**Appendix 1-Figure 1.**
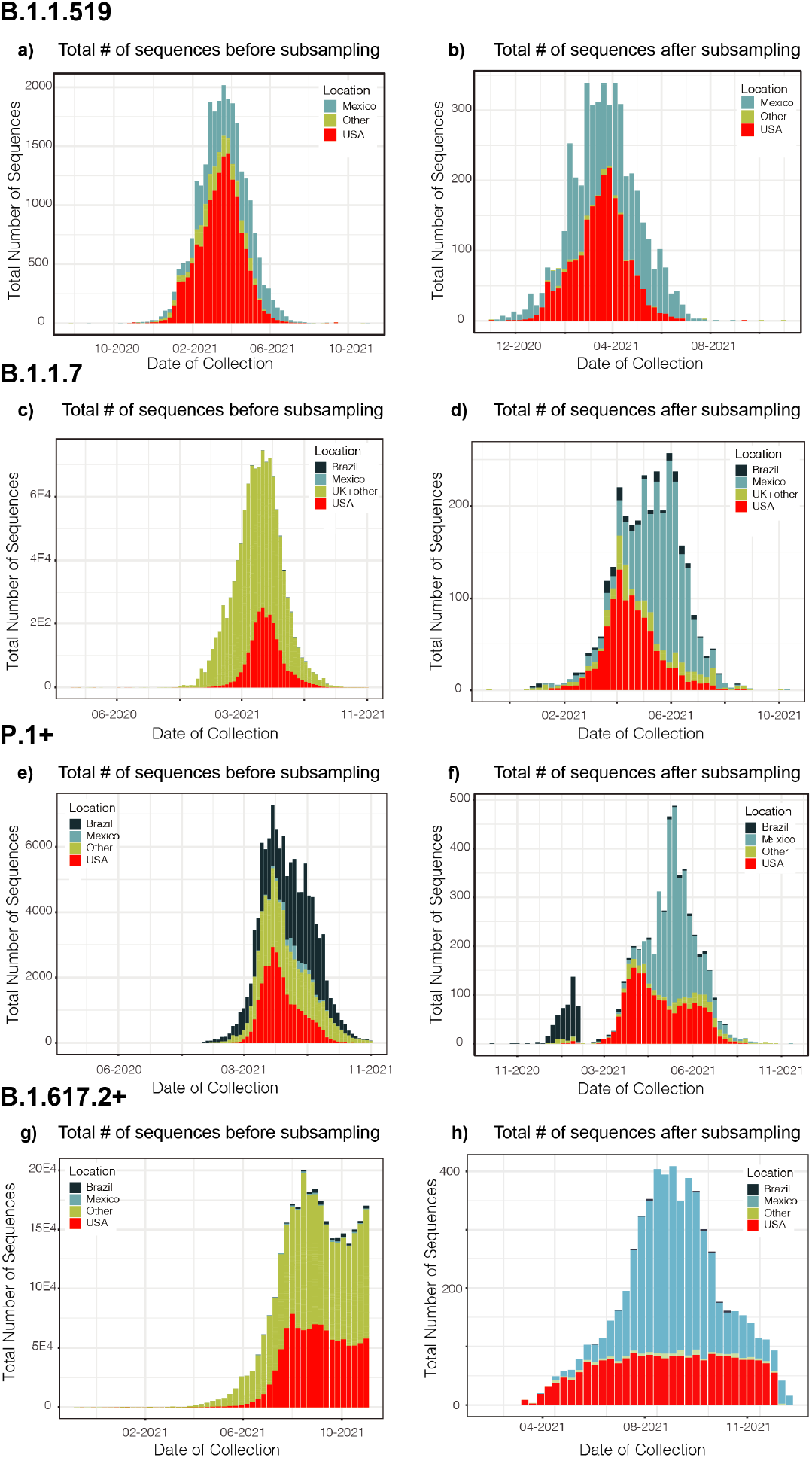
Distribution plots for each genome dataset before and after applying our migration- and phylogenetically-informed subsampling pipeline. Distribution plots for the number of genomes in the datasets before and after applying our subsampling pipeline. Plots for the B.1.1.519 (a and b), B.1.1.7 (c and d), P.1+ (e and f), and B.1.617.2+ (g and h) show the total number of sampled genomes coloured according to location, ranked according to the countries representing the most intense human mobility flow into Mexico derived from anonymized relative human mobility flow into different geographical regions.

**Appendix 1-Figure 2.**
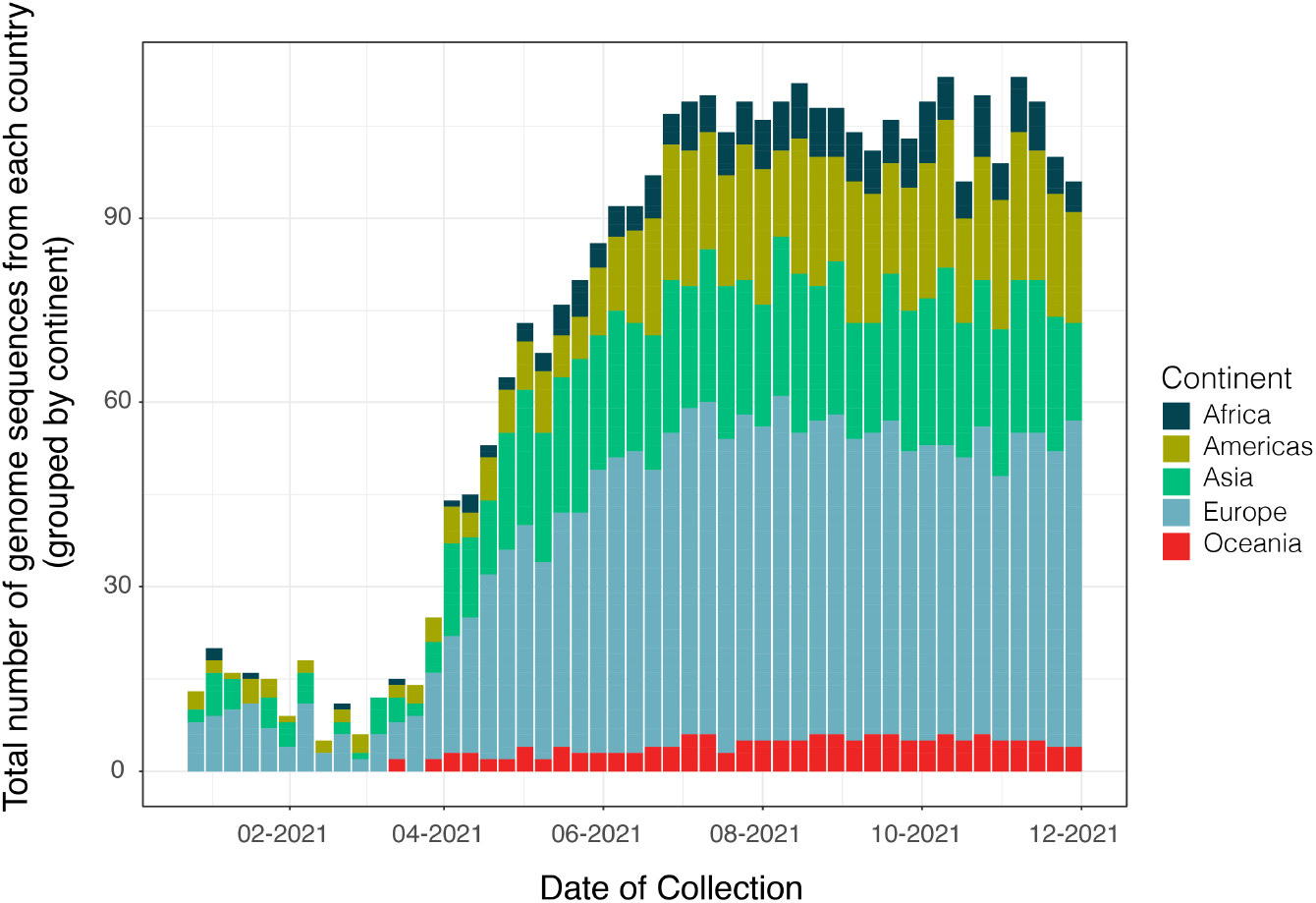
Distribution of genome sequences the new B.1.617.2+ dataset after subsampling under a different migration-informed approach (validation) Distribution of the number of genomes in the dataset corresponding to an alternative sub-sample of B.1.617.2+ sequences used for the validation of our migration informed subsampling approach. The dataset was built to obtain a homogeneous and proportional number of genome sequences from all countries sampled in GISAID (relative to their availability in the platform). The total number of genomes sequences sampled per region (represented by countries grouped by continent) are coloured according to their continent of origin. To compare to the distribution of genome sequences before subsampling, see **Appendix 1-Figure 1** above.

**Appendix 1-Figure 3.**
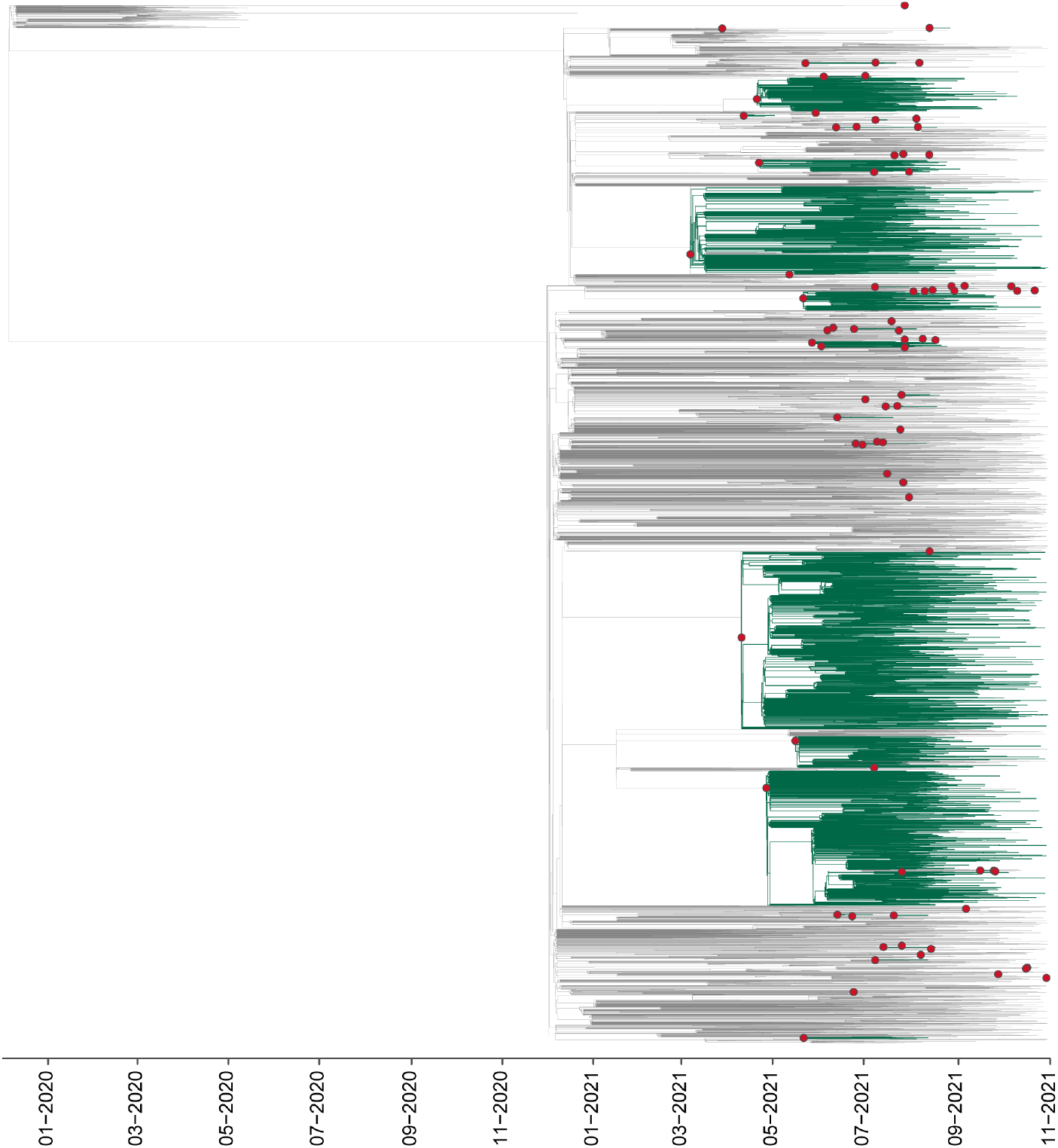
DTA analysis for the new B.1.617.2+ dataset (validation) Maximum clade credibility (MCC) tree for the alternative B.1.617.2+ dataset comprising a sub-sampling from all countries, represented by B.1.617.2+ sequences deposited in GISAID available up to November 30^th^ 2021, in which major clades identified as distinct introduction events into Mexico are highlighted. Nodes shown as red circles correspond to the inferred most recent common ancestor (MRCA) for clades representing independent introduction events into Mexico.

## Notes

https://github.com/rhysinward/Mexico_subsampling

### Competing Interest Statement

The authors have declared no competing interest.

### Summary of Updates

The main text has now been modified to frame the study within the hypothesis and goals stated. We have further aimed to connect key conclusions to public health implications, applied to the country and the extended geographic region of the Americas. We now include a new figure (Supplementary Figure 1 c) comparing the overall cumulative proportion of genomes generated per state between 2020-2021. Furthermore, we have also added additional maps representing the geographic distribution of the clades identified within Figures 2-4. The main text has been modified to provide further details on the sampling scheme, and we thoroughly discuss how this may impact our findings. We now provide a comparative analysis to validate our migration-informed genome subsampling approach. For this, we re-ran the time-scaled analysis applied to the B.1.617.2 dataset (representing the best sampled lineage in the country) using a different migration-informed sub-sampling scheme (including all countries). We then re-estimated the number of introduction events and compared the obtained value with our initial result. This is now presented in a new section in SI (Supplementary Text 2 and Figure 2). For the new dataset, a significantly lower number of introduction events into Mexico were inferred whilst C5d displayed a reduced diversity, further supporting for our migration-informed subsampling approach.

